# Circuit-specific selective vulnerability in the DMN persists in the face of widespread amyloid burden

**DOI:** 10.1101/2022.11.14.516510

**Authors:** Samuel J. Brunwasser, James N. McGregor, Kiran Bhaskaran-Nair, Clayton Farris, Halla Elmore, Eva L. Dyer, John M. Grady, Jennifer D. Whitesell, Julie A. Harris, Keith B. Hengen

**Author notes:** Equal Contribution (Samuel J. Brunwasser, James N. McGregor).

## Abstract

The relationship between brainwide functional decline and accumulation of pathological protein aggregates in Alzheimer’s disease (AD) is complex and not well understood. A set of highly interconnected cortical regions known as the default mode network (DMN) exhibits selective vulnerability to both functional decline and amyloid beta (A*β*) plaques in early AD. One possibility is that early A*β* accumulation in the DMN drives vulnerability. However, it is unknown whether there is something intrinsic to neuronal projections within the DMN that biases these circuits towards dysfunction. Here we directly test this hypothesis using long-term recordings of the spiking activity of ensembles of single units in freely behaving mice characterized by global cortical and hippocampal A*β* burden (APP/PS1). Specifically, we track the interactions of a population of neurons within a DMN region and two additional populations that comprise monosynaptic targets, one within and the other outside the DMN. In addition, we record single neurons in hippocampus and examine interactions between the in-DMN and out-DMN cortical circuits triggered on hippocampal sharp-wave ripples, stereotyped hippocampal events that contribute to memory consolidation in the cortex. We examine the statistics of local activity as well as inter-regional communication in a region, genotype, and brain-state dependent manner. Our data reveal dysfunction restricted to the in-DMN projecting circuit. In contrast, communication along neuronal projections that originate in the DMN but target an out-DMN population is equivalent in APP/PS1 and WT mice. Circuit dysfunction is most evident throughout sleep, and particularly disrupted within sharp-wave ripples. Summarily, our results indicate that, even in the face of transgene overexpression and widespread A*β*, there is distinct intrinsic and selective vulnerability. This vulnerability to amyloidosis is circuit-specific and conditioned on target, and neither source nor amyloid burden. These data raise the possibility that neuronal function in the DMN is not universally vulnerable; DMN subnetworks whose interactions involve targets outside the DMN may be resilient to A*β*.

## INTRODUCTION

The default mode network (DMN) is a set of distributed but highly interconnected brain regions that are coactivated during wakeful rest^1^. In addition to its role in cognition, the DMN is noteworthy as it is selectively vulnerable to the early stages of neurodegenerative disease. Specifically, in Alzheimer’s Disease (AD), amyloid-beta (A*β*) accumulation starts in DMN hubs and propagates to secondary regions along anatomical connections^2–4^. Unsurprisingly, brain regions that accumulate A*β* show disrupted activity^5^; disrupted functional connectivity between DMN regions is a hallmark of early AD^6^.

The simplest explanation is that changes in the DMN are purely and directly a result of A*β*. This rationale supports the prediction that A*β* deposits in other circuits would render them similarly disrupted. Alternatively, early AD represents the confluence of two events. Specific circuits are intrinsically vulnerable to A*β*-related damage, and it is in these circuits that A*β* takes hold. In this case, if A*β* deposits were widespread, DMN circuitry would still be disrupted relative to non DMN regions. However, because A*β* deposition is restricted to the DMN early in AD, it is difficult to separate the local impact of A*β* from an intrinsic vulnerability of these brain regions. Either possibility has powerful implications for an understanding of AD etiology and intervention strategies. Here we take advantage of a mouse model of widespread amyloidosis and long-term, multi-site recordings spanning in-DMN and out-DMN circuits to directly test the hypothesis that neurons in the DMN are intrinsically vulnerable independent of A*β*.

The regions comprising the DMN exhibit a high degree of reciprocal connectivity^7–9^, such that the majority of DMN efferents target other DMN regions. Nonetheless, a minority of DMN projection neurons target postsynaptic sites broadly throughout the brain^10^. In both cases, the cell bodies lie within the anatomical borders of the DMN, but the circuits are divisible by postsynaptic target; in-DMN projections originate and terminate in DMN regions, while out-DMN projections terminate in non-DMN regions. If DMN neurons are intrinsically vulnerable to A*β*-related dysregulation, the dissociation of DMN and non-DMN circuitry by projection target raises an interesting possibility: the cellular impact of disease may be conditioned on axonal target, even when comparing adjacent neurons.

The recent discovery of two neuronal populations in the mouse ventral retrosplenial cortex (RSPv), a core DMN region, reveals such intermingled circuitry^10^. Both subpopulations are composed of excitatory projection neurons, but differ in their postsynaptic targets. One group, the in-DMN targeting, sends monosynaptic connections to dorsal anterior cingulate (ACAd), another selectively vulnerable DMN region. The other group, the out-DMN targeting, sends monosynaptic connections to primary visual cortex (VISp), which is outside of the DMN. This anatomy crystalizes the question of selective vulnerability, allowing for interrogation not only of region but also of connectivity. For three reasons, this circuitry, in combination with the APP/PS1 mouse, is ideally suited to test the hypothesis that subsets of neurons are intrinsically vulnerable during disease independent of the anatomical pattern of A*β* deposition.

First, the APP/PS1 mouse overexpresses amyloid precursor protein (APP) and presenilin globally and is remarkable for indiscriminate A*β* accumulation throughout the cortex and hippocampus^11^. This deposition pattern thus circumvents the confounding restriction of A*β* to the DMN. Second, mouse models are amenable to multi-site recording of ensembles of single neurons, allowing for the simultaneous monitoring of multiple monosynaptically connected populations in the context of freely behaving animals. Finally, the APP/PS1 mouse is generally accepted as a model of early AD progression because disruptions in cortical physiology are not explained by neuron loss^12^.

Here we collected and analyzed four-site recordings of neuronal ensembles in APP/PS1 mice to test the hypothesis that two circuits, one in-DMN (RSPv-to-ACAd: R2A) and one out-DMN (RSPv-to-VISp: R2V), are differentially vulnerable in the context of APP overexpression, PS1 overexpression, and global amyloid burden (APP/A*β*). We conducted long-term, simultaneous recordings of extracellular spiking from ensembles of single neurons in RSPv, ACAd, VISp, and hippocampus (CA1) of freely behaving adult WT and APP/PS1 mice. We examined neuronal dynamics and circuit interactions as a function of brain state over a 24 h cycle containing a full range of naturally occurring behaviors. Our results reveal selective functional disruption only in the R2A circuit, and this effect is pronounced during sleep. Crucially, both the R2A and R2V circuits originated from the same population of neurons in the RSPv. These data suggest that, even within the same DMN hub region and global expression of APP/A*β*, intrinsic vulnerability is determined by projection target.

## RESULTS

### Chronic Monitoring of Single Neuron Activity in Four Brain Regions in WT and APP/PS1 Mice

Mice harboring the APP/PS1 transgenes exhibit increased brain A*β* by 8 months and neuritic plaques in the isocortex by 10 months of age^13^. While A*β* expression is constrained to the DMN in the earliest stages of human AD, we observed plaques throughout isocortex and hippocampus of APP/PS1 animals (Fig. S1A,B) as described previously^11^. To test the hypothesis that circuits whose source and target are within the DMN are intrinsically vulnerable in the context of APP/A*β*, we performed continuous (10-20 d), multisite recordings of extracellular signals from ACAd, RSPv, VISp, and CA1 (Fig.1A-B) in APP/PS1 and WT littermates (n = 4 WT age 13.6 +/− 0.85 months, n = 4 APP/PS1 age 13.8 +/− 1.45 months, see Table 1 for animal-by-animal details). (NB a circuit in this context refers to the subset of neurons within a given region whose activity is relayed via projections to a target region. Both local/recurrent and projecting neurons are encapsulated by the term circuit.)

**Figure 1:**
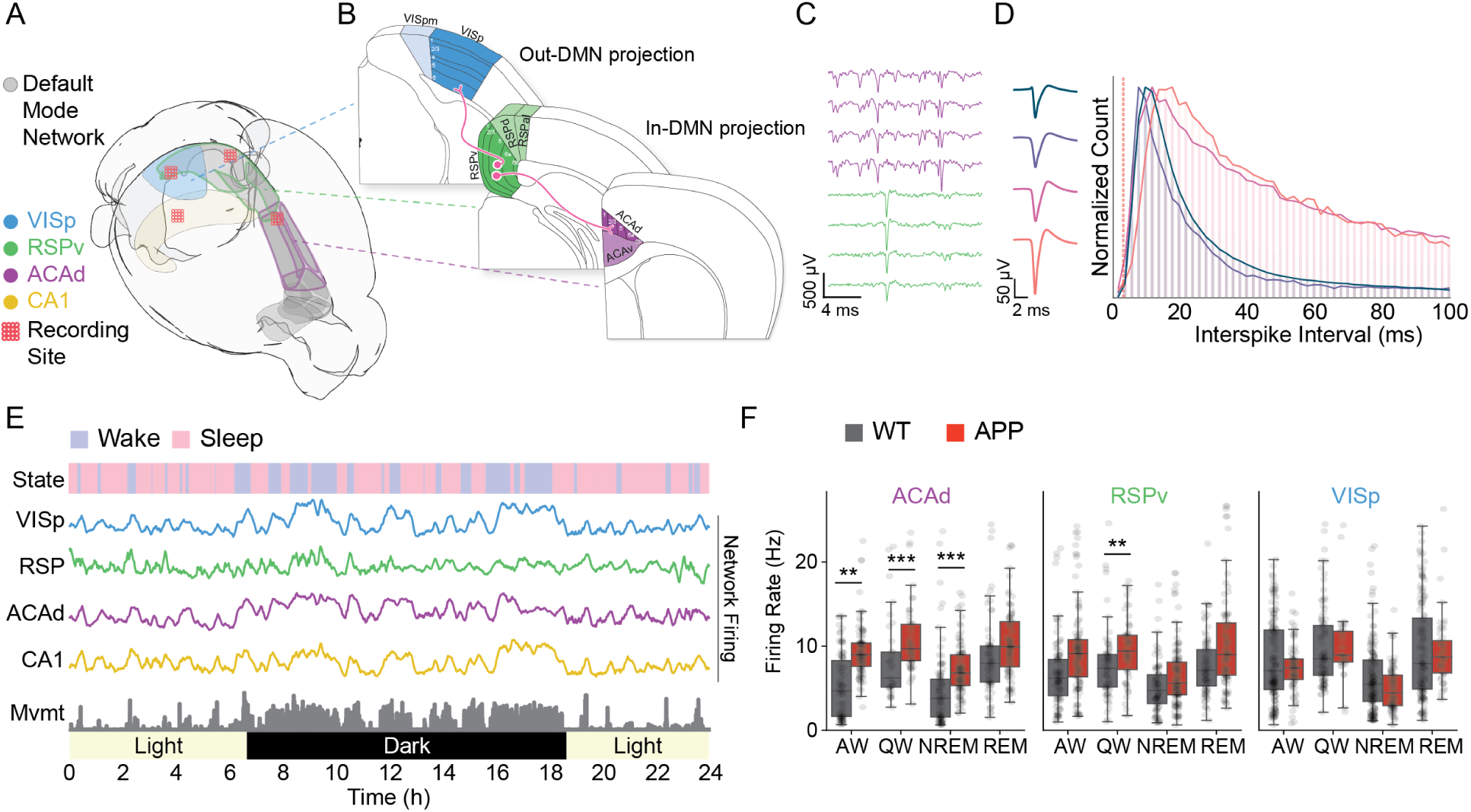
A*β* overexpression drives increased neuronal activity in DMN regions as a function of brain state. **A.** Experimental overview. 64 ch tetrode bundles were implanted in four brain regions (total of 256 ch), two in the default mode network (DMN; RSPv, ACAd), one outside of the DMN (VISp), as well as CA1 hippocampus. **B.** Illustration of the monosynaptic projection connecting populations in RSPv to ACAd (in-DMN) and population in RSPv to VISp (out-DMN). **C.** Example raw data trace from two tetrodes, one in RSPv and the other in ACAd. **D.** (left) four mean waveforms of single units recorded on one tetrode and their (right) interspike interval histograms which reveal a refractory period. Vertical dashed red line indicates minimum possible refractory period for a single unit. Colors in the histogram correspond to the 4 unit waveforms on the left. **E.** 24 h of data from a single animal. Rows denote sleep/wake states, mean ensemble firing rate by region, animal locomotion (“movement”), and environmental conditions (light/dark). **F.** Mean firing rate of all single units as a function of genotype, region, and brain state (* p *<* 0.05,*** p *<* 0.001; linear mixed model log(firing rate) genotype * state * region + (1—animal) + (1—epoch) + (1—neuron), where animal, epoch, and neuron are included as random effects). VISp: primary visual cortex; RSPv: ventral retrosplenial cortex; ACAd: dorsal anterior cingulate cortex; CA1: cornu ammonis hippocampus; WT: wild-type; APP: amyloid precursor protein and mutant human presenilin 1; AW: active wake; QW: quiet wake; NREM: non-rapid eye movement sleep; REM: rapid eye movement sleep.

**Table 1:** Animal demographics and implantation timing.

| ANIMAL | GENOTYPE | SEX | AGE AT IMPLANTATION |  |
| --- | --- | --- | --- | --- |
|  |  |  | PND | MONTHS |
| CAF69 | WT | F | 374 | 12.5 |
| CAF71 | APPS1 | M | 488 | 16.2 |
| CAF73 | APPS1 | F | 374 | 12.5 |
| CAF74 | APPS1 | M | 412 | 13.7 |
| CAF75 | APPS1 | M | 383 | 12.8 |
| CAF77 | WT | F | 393 | 13.1 |
| CAF81 | WT | F | 432 | 14.4 |
| CAF82 | WT | M | 435 | 14.5 |

**Table 2:** Recording data and epoch counts across sleep-wake states.

| ANIMAL | REC.<br>TIME<br>(HR) | # UNITS<br>RECORDED | # EPOCHS |  |  |  |
| --- | --- | --- | --- | --- | --- | --- |
|  |  |  | AW | QW | NREM | REM |
| CAF69 | 35.75 | 156 | 165 | 161 | 58 | 40 |
| CAF71 | 20.84 | 121 | 180 | 235 | 104 | 7 |
| CAF73 | 24 | 216 | 113 | 112 | 49 | 16 |
| CAF74 | 24 | 163 | 192 | 281 | 98 | 5 |
| CAF75 | 24 | 205 | 242 | 321 | 85 | 6 |
| CAF77 | 23.84 | 58 | 176 | 206 | 78 | 34 |
| CAF81 | 24 | 228 | 269 | 300 | 69 | 43 |
| CAF82 | 24 | 112 | 243 | 259 | 69 | 54 |
| **TOTAL** | **200.43** | **1259** | **1580** | **1875** | **610** | **151** |

Single units were extracted with MountainSort-based clustering^14^ and labeled automatically by an XG Boosted decision tree^15^ trained on measures of cluster quality and isolation such as L-ratio, Mahalanobis distance, and the absolute refractory period (Fig.1C,D). Regular spiking units (RSUs, ∼ 90% pyramidal) were classified based on waveform^16–18^. In addition, high frame-rate video was collected and processed with a convolutional neural network to track animal locomotion^19^. For sleep scoring, this was combined with 60 Hz low-passed electrophysiological data as described previously^20^ (Fig. S1F-G). Here we analyze 24 h of data from each animal, collected at least 1 w into the long-term recording, and three weeks after surgery. Analyses are restricted to well-isolated RSUs with a presence ratio of at least 80% (i.e., included neurons exhibited detectable activity for the entire period of analysis). Each animal’s RSPv ensembles contained single units with short-latency interactions (significant single neuron pairwise correlations at *<* 10 msec) within both ACAd and VISp, consistent with monosynaptic excitatory connectivity (Fig. S3B,C). These results are consistent with monosynaptic projections from RSPv to both ACAd and VISp, as described recently in a study of mesoscale connectomics^10^.

### Hyperexcitability is restricted to DMN circuitry in APP/PS1 mice and progresses with age

There are two key features of amyloidosis driven by APP overexpression that we sought to replicate to establish consistency between our recordings and the literature. First, it is difficult to disentangle the effects of A*β* accumulation from the effects of constitutive overexpression of APP and PS1. A key difference is that A*β* accumulation is progressive, such that its downstream effects change with age^21,22^. Second, amyloidosis (and APP overexpression) disrupt neuronal firing rates, albeit with conflicting evidence of direction: calcium transients across frontal cortex and CA1 suggest hyperexcitability^23,24^, while extracellular recordings in frontal cortex indicate hypoexcitability^25^. Experimental constraints, such as head-fixation and relatively short recordings, make it difficult to reconcile these findings. To address each of these points, we examined single neuron activity as a function of brain region, arousal state, and genotype. Crucially, our approach of continuous, long-term recording allows us to follow the activity of the same single units across behavioral and environmental conditions. As a result, a linear mixed model (LMER^26^) can evaluate the interaction of circuit, state, and genotype while quantifying variance explained by individual animals as well as individual neurons. We also took advantage of the fact that, despite all of our animals being *>* 1 y, there was a 114 d range between the youngest and oldest animal in our dataset (Table 1). The resultant linear mixed model was log(firing rate) ∼ genotype × state × region × age + (1|animal) + (1|cell ID), where animal ID and cell ID are included as a random effect.

Broadly, there were no main effects of either genotype (*p* = 0.823) or age (0.397). However, the three way interaction between age, genotype, and region was significant (*p* = 5.708*e* − 15). To explore this interaction, we conducted post-hoc tests using EMMeans (R). Consistent with prior evidence of hyperexcitability, firing rates were significantly higher in the ACAd and RSPv of APP/PS1 compared to WT (*p* = 0.038 and 0.016, respectively). Despite near-global plaque coverage (Fig. S1A,B), this effect was circuit specific: there was no difference between APP/PS1 and WT firing rates in CA1 or VISp (*p* = 0.501 and 0.166, respectively).

Additionally, there was a significant interaction between genotype, brain state, and brain region (*p <* 2.2*e* − 16). This was driven by state-dependent differences between WT and APP in both ACAd and RSPv, but neither VISp nor CA1 (Fig. 1F; significant comparisons between genotypes: RSPv quiet wake, *p* = 0.0026; ACAd quiet wake, *p* = 0.0008; ACAd NREM, *p* = 0.0004; ACAd active wake, *p* = 0.0011. Results in RSPv during active wake and REM had *p* = 0.099 and 0.094, respectively. Collectively, altered firing rates are unlikely to arise from gross A*β*-related damage; single neuron yield did not differ by genotype (Fig. S1C,D), consistent with a lack of cell loss at 1 y. Neuron counts in representative histology were similar in each recorded region (Fig. S1C). Taken together, these results suggest that neuronal activity levels in RSPv and ACAd are selectively sensitive to APP/A*β*, in contrast to VISp and CA1.

The model above suggests that age explains a significant amount of variance in the firing rate of neurons in RSPv and dACC of APP/PS1 mice. For a simple and direct comparison of the impact of age as a continuous variable, we made separate models for APP/PS1 and WT animals log(firing rate) ∼ state × region × age + (1|animal) + (1|cell ID), where animal ID and cell ID are included as random effects but only including animals from a given genotype). In the WT model, the main effect of age was not significant (*p* = 0.712), while in the APP/PS1 model, it was (*p* = 0.001). Grouped together, in the APP/PS1 model, the DMN regions (RSPv and ACAd) exhibited significantly higher firing rates than non-DMN regions (VISp and CA1) (*p* = 0.041). In the WT model, the opposite was true: non-DMN regions exhibited higher firing rates than DMN regions (*p <* 0.001).

Summarily, even after 1 y of age, the effect of genotype on firing rate is progressive, but only in the ACAd and RSPv, and only in APP/PS1 animals. These effects are consistent with prior descriptions of A*β*^22^, but do not rule out significant contributions from the overexpression of APP and PS1.

### Timing within the R2A but not R2V circuit is disrupted in APP/PS1 mice

While ensemble mean firing rates were elevated in a state-specific fashion in the two DMN regions, these circuits are complex and contain information that projects both locally and to diverse post-synaptic targets, some of which are contained within the DMN and some of which lie outside of the DMN. Ensemble mean firing rates do not provide insight into these distinct computational processes.

It is possible that functionally distinct yet anatomically intertwined subcircuits are differentially vulnerable to disease. We reasoned that we could gain insight into the integrity of functional subsets of recorded regions by examining the strength and timing of population-level interactions. For example, neurons within RSPv process and then send information along monosynaptic projections to ACAd. Neurons in ACAd receive this transmission and, through additional local interactions, generate stereotyped responses. As a result, the activity of a subset of neurons in RSPv should be related to the activity of a subset of neurons in ACAd within low latency (tens of ms), even if they (the RSPv neurons) are not projection neurons. Conceptually, this is similar to studying lagged functional connectivity in imaging or LFP data, albeit with single unit resolution^27,28^. In this context, alterations in population-level interactions could arise from a variety of mechanisms, such as disruption of local interneuron timing or degradation of anatomical lines. Thus, studying population interactions with single-neuron resolution is well positioned to capture the summary effect of disrupted communication between brain regions.

Multi-site recordings allow us to ask whether the vulnerability of RSPv, a DMN hub, is a general feature of the region, or if it is specific to functional sub-circuits. To address this, we performed lagged pairwise spike correlation between all pairs of single units in the RSPv (source region) and each of two target regions (ACAd and VISp) (Fig. 2A). We analyzed these population-wide interactions as a function of each epoch of NREM and waking. Crucially, the same population (RSPv) serves as the source region in each of these circuits. We measured the magnitude and directionality (delay) of the source/target interaction as a function of genotype and arousal state (420 wake epochs, 832 NREM epochs) (Fig. 2B,C).

**Figure 2:**
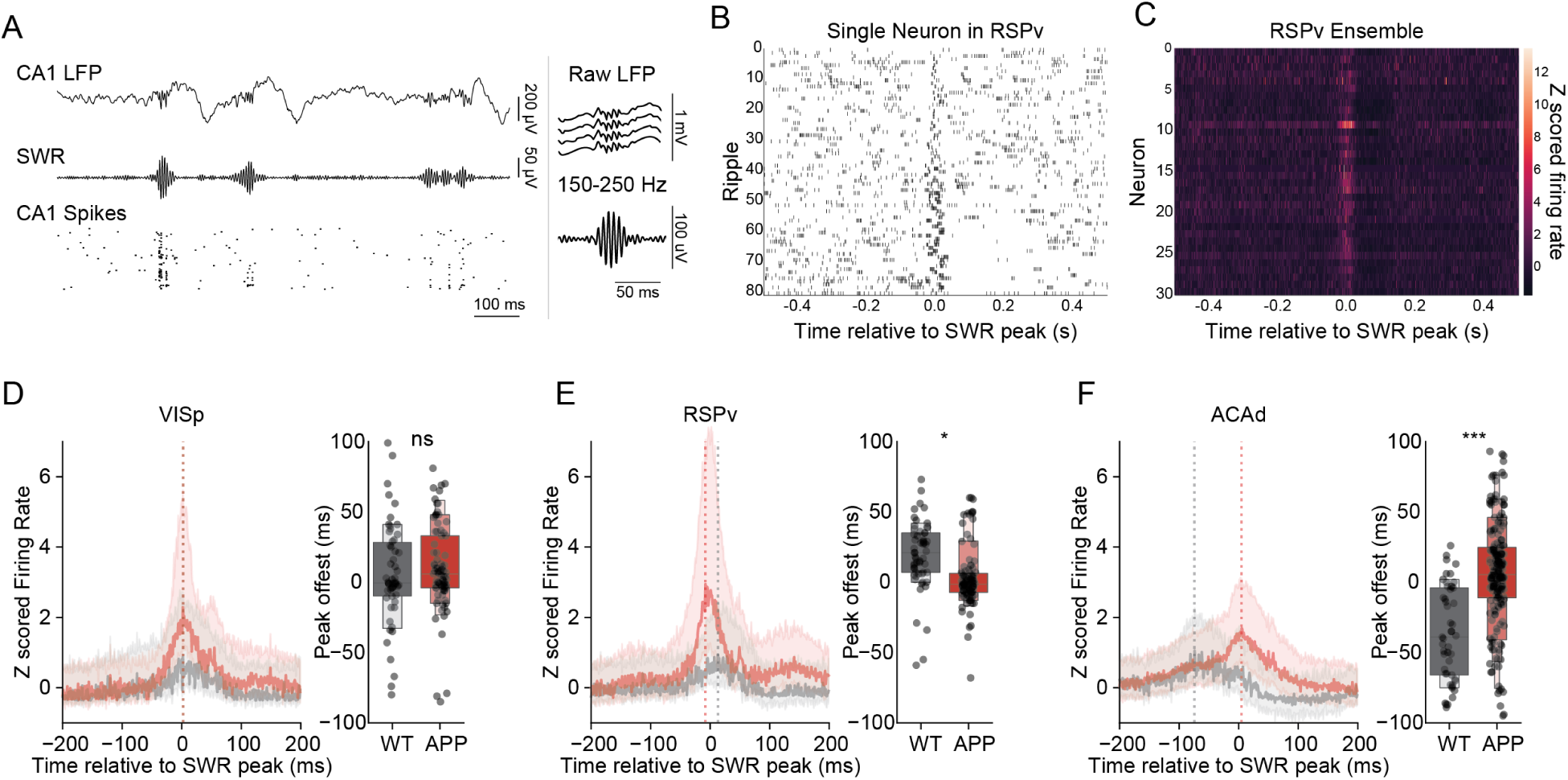
The timing of NREM sharp-wave ripple-driven activity is disrupted only in the DMN. **A.** Cartoon illustrating source/target lagged correlation analysis. In each behavioral epoch, single neuron activity in RSPv is correlated with single neuron activity in each of two targets. Epochs of the same type are then averaged together. **B.** The timing of R2V synchrony does not vary as a function of genotype or brain state. **C.** Quantification of the time difference between source and target peaks. There are no significant differences between genotypes in out-DMN circuit timing in either wake or NREM sleep. **D.** The timing of R2A synchrony does vary as a function of genotype and brain state. **E.** Quantification of time difference between RSPv and ACAd peaks. Relative to WT, APP target activity is delayed significantly in NREM but not wake (WAKE: p = 0.173. NREM: p = 0.011, linear mixed effects model: peak timing genotype * state * circuit + (animal) where animal is a random effect). **F.** The timing of R2A synchrony varies as a function of active vs. quiet wake. **G.** Quantification of time difference between RSPv and ACAd peaks. APP target activity is significantly delayed compared to WT in quiet wake but not active wake (AW: p = 0.243. QW: p = 0.01, linear mixed effects model: peak timing genotype * state * circuit + (animal) where animal is a random effect).

Consistent with prior reports of broadly increased functional connectivity relative to WT mice^29^, both R2A and R2V circuits in APP/PS1 animals exhibited an increased correlation magnitude relative to WT (Fig. 2B,C, S4A,B; genotype/circuit interaction p = 3.716e-10, linear mixed model where animal ID was treated as a random effect). There was no significant effect of state in modulating the increased correlation magnitude (p = 0.262).

Within this context, we examined the temporal dynamics of inter-regional communication. When considering timing, the three-way interaction of brain state, genotype, and circuit was significant (p = 0.019). In APP animals, the RSPv to ACAd (R2A) interaction was lagged by 6 msec compared to WT animals, an effect only observed during NREM sleep (p = 0.011). In contrast, in the same animals, RSPv to VISp (R2V) interaction timing was not different from WT in either state (wake: p = 0.732; NREM: p = 0.548). The same analyses run on shuffled spike times yielded flat lines with no peaks (Fig. S4C-F), supporting that these analyses capture biologically meaningful structure.

Although we did not observe a significant difference in the timing of R2A interaction between genotypes during wake, meaningful differences in neuromodulatory tone and neural dynamics exist between active waking (AW) and quiescence (QW). Analyzed as waking substates, the R2A interaction in APP mice was lagged by 5 msec compared to WT animals during QW (p = 0.01), but not during AW (p = 0.243) (Fig. 2F,G). The timing of R2V interaction did not significantly differ between genotypes in either AW or QW (AW: p = 0.874; QW: p = 0.297).

Because each circuit shared the same source neurons, these results suggest that selective vulnerability in APP/A*β* may not simply reflect anatomical proximity to plaques, but instead depends on circuit details, such as connectivity. Further, given the state-specificity of these results, our data suggest that disruptions in communication are unlikely to be the result of amyloid-driven anatomical damage, as this would be expected to manifest independently of state. A plausible hypothesis is that amyloid plaques alter the impact of neuromodulators on a subset of cortical interneurons that are over-represented in the processing and transmission of information along a specific set of projections, such as in-DMN circuits^30–32^.

Epochs of NREM can last for thousands of seconds and contain diverse substates. Collapsing across this heterogeneity reveals state- and circuit-specific effects of APP/A*β*, but does not provide insight into the possible disruption of functionally-relevant neurophysiological events within NREM sleep.

### The timing of sharp-wave ripple-driven spiking in RSPv is aberrant in APP/PS1 animals

We reasoned that if altered communication is specific to the R2A circuit, neurophysiological events known to drive fluctuations in the DMN might be particularly vulnerable to the impact of APP/A*β*. An ideal candidate for such an event is the sharp-wave ripple (SWR), which is a brief large deflection and oscillation (140-220 Hz) in the local field potential of the hippocampus that has been linked to memory consolidation between the hippocampus and cortex^33,34^. Notably, SWRs drive subsequent DMN activity^35^. We next asked whether SWRs are consistent with a locus of selective vulnerability in the R2A circuit.

Arising from the dendritic layer of CA1, SWRs are observed primarily during NREM sleep but also quiet waking. The discrete, aperiodically repeating nature of SWRs presents an opportunity to approach long term recordings of freely behaving animals with precise, trial-structured analysis. To understand intra-regional communication in this context, we first identified ripples in broadband CA1 data during NREM epochs using standard algorithms^36^ (Fig. 3A). We observed no significant effect of genotype on SWR peak frequency, amplitude, or duration, although there was noteworthy inter-animal variability in all three measures, particularly in WT animals (Zhurakovskaya et al., 2019^37^, but see Cushing et al., 2020^38^) (Fig. S5A-D, Wilcoxon rank sums ripple density, duration, amplitude, and peak frequency p *>* 0.05). While SWRs are generated in the hippocampus, one of the major inputs to RSP is a monosynaptic projection from CA1^39^, and SWR driven spiking in RSP has been described previously^40^. In our data, individual neurons in RSP displayed increased firing around the time window of hippocampal SWRs (Fig. 3B,C).

**Figure 3:**
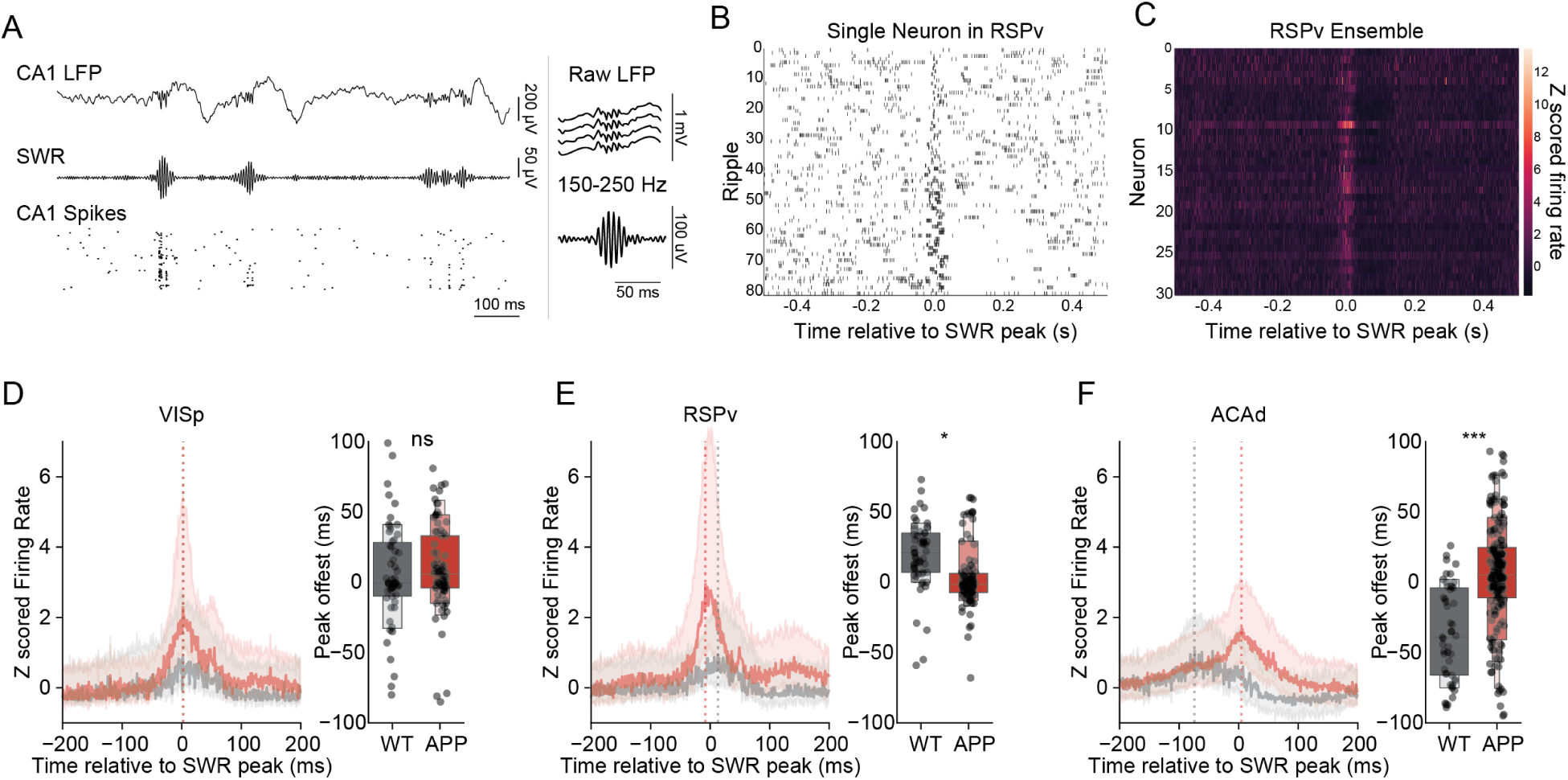
Inter-regional spike timing during sleep is disrupted in the R2A circuit. **A.** (left) Example sharp wave ripples (SWRs) shown in: (top) raw local field potential (LFP) recorded in CA1, (middle) 150-250 Hz band-passed data, and (bottom) CA1 single unit activity. (right, top) A single SWR in raw data, and (right, bottom) band-passed data. **B.** Raster plot of the response of a RSPv single unit to 80 SWRs. **C.** The average response of 30 single units to SWRs. **D.** The timing of single unit activity during NREM SWRs in VISp does not differ by genotype. (left) Peri-ripple histogram across units from all mice, (right) distributions of peaks by genotype (linear mixed effects regression: peak timing genotype * region + (animal), where animal is a random effect) **E.** RSPv single unit activity during NREM SWRs is significantly accelerated compared to WT. **F.** ACAd single unit activity during NREM SWRs is significantly delayed relative to WT. n.s. not significant, *** p *<* 0.001.

We reasoned that, in the context of general sleep and wake, R2A circuitry was delineated from R2V circuitry by perturbation of the timing of intraregional communication. The hippocampus is sometimes considered part of the DMN^41^, thus we hypothesized that the timing of SWR activation of RSPv and ACAd (i.e., DMN regions) but not VISp (non-DMN) should be aberrant in APP/PS1 animals relative to WT. In line with our hypothesis, when examining SWR timing, there was a significant interaction between genotype and circuit (*p <* 2*e* − 16, linear mixed model where animal ID was treated as a random effect). Interestingly, we only saw timing disruptions in R2A circuitry. The timing of SWR-related activity in VISp did not differ by genotype (Fig. 3D; Median peak time WT 2 +/− 4.9 ms, APP 2 +/− 3.3 ms, p = 0.161). However, in APP/PS1 animals, the SWR peak in RSPv was accelerated (Fig. 3E; median peak time WT 13 +/− 3.2 ms, median peak time APP −8 +/− 2.2 ms, p = 0.021). In addition, SWR-related spiking in RSPv of APP/PS1 animals was characterized by a second, slower peak that was not observed in WT animals. Further, we observed a significant delay in the timing of SWR-associated spiking in ACAd (Fig. 3F; median peak time WT −74 +/− 4.9 ms, APP 5 +/− 3.3 ms, p *<* 0.0001).

Consistent with our observations of a global increase in pairwise correlations, the response magnitude of RSPv neurons in APP/PS1 animals to SWRs was exaggerated relative to WT (Fig. 3E; peak height p = 1.24e-8; median peak height WT = 4.52, APP = 6.50). This effect was similar in VISp but to a lesser extent (Fig. S5E-G). These responses were not observed outside of SWRs in either genotype (Fig. S6). In addition, while responses to SWRs during wake were observed in all three regions, they were smaller in magnitude than in NREM and did not differ by genotype (Fig. S7). We also examined responses to NREM SWRs in fast spiking interneurons and noted increased responses in all regions in APP mice without significant changes in timing (Fig. S8).

Taken together, these results suggest that APP/A*β* enhances the response of local populations of putative excitatory neurons to SWRs in the RSPv, and that the timing of local SWR-driven computation is disrupted in the R2A ensembles but not the R2V ensemble.

### APP/A*β* drives altered communication dimensionality in a circuit- and state-dependent fashion

While these data suggest that hippocampal projections to DMN regions are intrinsically vulnerable to APP/A*β*, it is unclear whether altered timing implies a disruption in the information content carried along a circuit. It is possible that, despite being delayed or accelerated by a few milliseconds, information transmission is otherwise normal. In this case, it is unlikely that minor disruptions in timing would represent biologically significant changes.

To assess the impact of APP/A*β* on the content of inter-regional communication, we measured the dimensionality of the communication subspace between each pair of regions. To do this, we performed reduced rank regression between source and target regions, first in the R2A and R2V circuits^42^. Regression performance, or the degree to which source activity predicted target activity, was measured as a function of the rank of the coefficient matrix (i.e., the number of predictive dimensions). When comparing genotypes this way, a consistent shift in the number of source dimensions necessary to capture a given percentage of the target activity indicates a change in the dimensionality of the interaction of the two regions (Fig. 4A). In simple terms, complex patterns of activity require more dimensions to reproduce than simple patterns. Applied to data such as ours, this analysis measures subspace dimensionality^42^: a subspace of the activity in RSPv predicts activity in VISp, and another subspace of RSPv activity predicts ACAd activity. It is important to note that subspaces are latent patterns of activity that often call on overlapping sets of individual cells, both local and projecting neurons.

**Figure 4:**
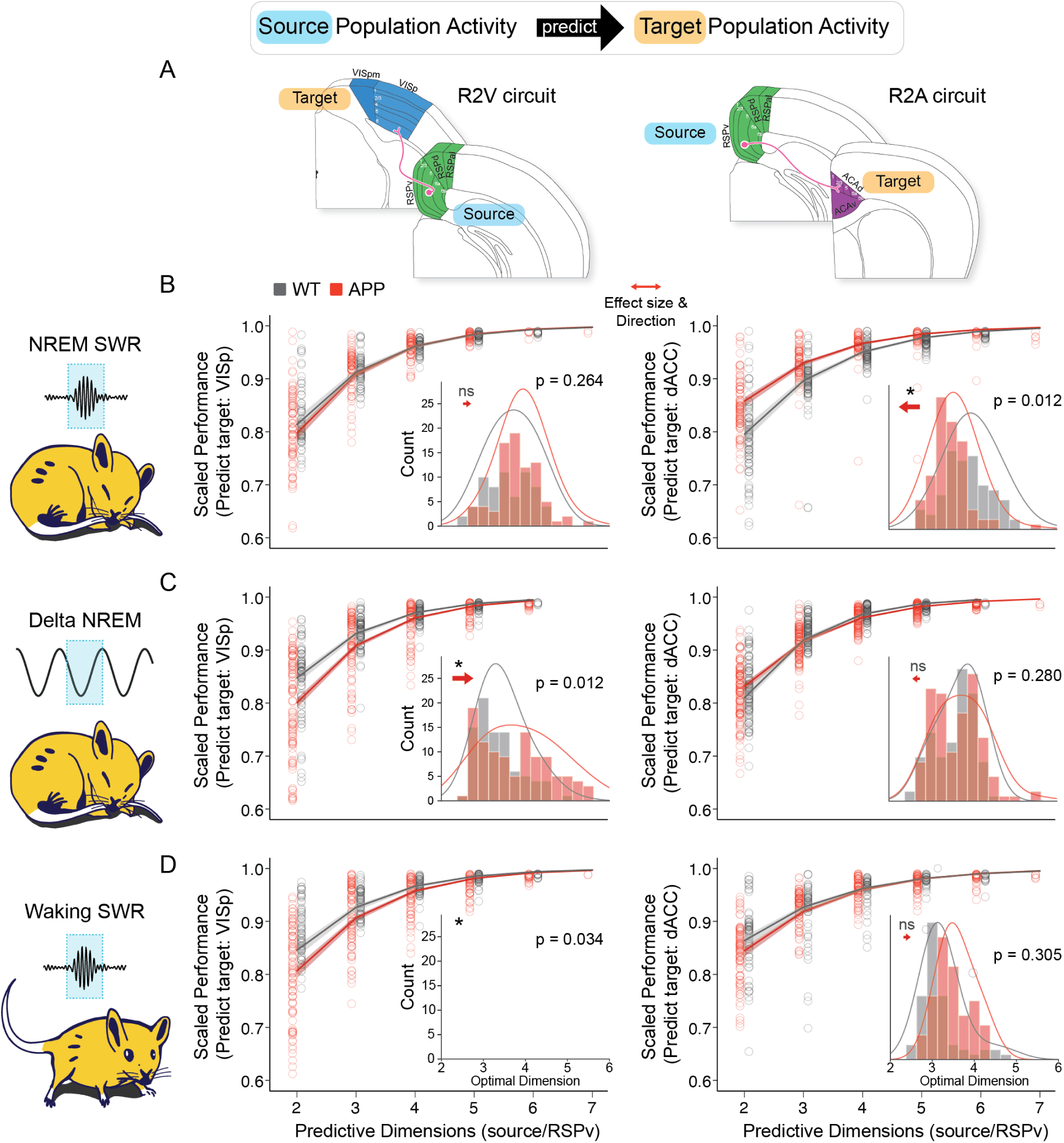
Communication in the R2A circuit exhibits reduced dimensionality during NREM sharp wave ripples but not other states. **A.** (top) Conceptual diagram of analysis. The activity of an ensemble of single neurons in a source region is used to predict the activity of neurons in the target region. The dimensionality of the source ensemble is then progressively reduced, and the loss of predictive power is measured. (bottom) Diagram of the R2A circuit and R2V. In each case, the source is the same: the RSPv. **B.** Dimensionality of R2V and R2A circuits in NREM sharp wave ripples (SWRs) as a function of genotype. Cartoon illustrates the behavioral/neurophysiological state. The left data column is from R2V, the right column is R2A. As the dimensionality of the source data is experimentally reduced, the predictive performance (scaled to proportion of the full prediction) is plotted. Each point represents one bootstrap, and the fitted line is a mixed model regression fit (logit(scaled performance) genotype * circuit * state + (animal/date/epoch/bootstrap). The last term represents nested random effects, see methods). The inset shows distribution of the optimal dimension (defined as the number of dimensions required to explain 90% of the full interaction) by genotype. The red arrow shows the direction of the effect (WT-APP) and the effect size (length of arrow). **C.** Same as B but during randomly identified, slow wave (delta: 0.1-4 Hz) enriched, non-SWR intervals of NREM sleep. **D**. Same as B but in SWRs occurring during waking (* = p *<* 0.05, ns = not significant).

We first examined subspace dimensionality during NREM SWRs (Fig. 4A-B). SWRs are specific events that drive activity in all recorded regions and thus provide trial-like structure in the context of free behavior. Despite the fact that SWR-driven spiking was increased in both VISp and RSPv in APP/PS1 mice relative to WT (Fig. S5), the relationship between the number of source dimensions and the predictability of target activity did not differ by genotype in the R2V circuit (RSPv to VISp). In this context, we defined the optimal dimensionality of circuit interactions as the number of dimensions necessary to capture 90% of the full rank prediction (Fig. 4B, left; p = 0.26, effect size = 0.23 where effect size is defined as WT optimal dimensionality minus APP optimal dimensionality. Linear mixed model regression: optimal dimension genotype * circuit * cond + animal/date/epoch/bootstrap. The last term represents nested random effects with random intercepts). In contrast, optimal dimensionality was significantly reduced during SWRs in the R2A circuit (RSPv to dACC) in APP/PS1 animals relative to WT (Fig. 4B, right; p = 0.012, effect size = −0.54). Succinctly, the complexity of circuit communication during NREM SWRs was reduced in APP animals, but only in the R2A circuit.

One possibility is that the reduction of dimensionality in the R2A circuit is general and not specific to NREM SWRs. To test this, we ran the same analysis in two additional contexts: 1) centered on periods of NREM sleep outside of SWRs (i.e., “delta NREM”, times enriched in slow wave activity), and 2) centered on SWRs occurring during waking^34^. During delta NREM intervals, we observed a reversal of the pattern described above; optimal dimensionality was increased in the R2V circuit of APP/PS1 animals (Fig. 4C, left; p = 0.012, effect size = 0.54). In the R2A circuit, there was no significant difference between APP and WT optimal dimensionality (Fig. 4C, right; p = 0.28, effect size = −0.21). Remarkably, this pattern was maintained in waking SWRs: dimensionality in the R2V circuit was significantly increased in APP animals (Fig. 4D, left; p = 0.034, effect size = 0.45), and there was no significant difference between genotypes in the R2A dimensionality (Fig. 4D, right; p = 0.305, effect size 0.207). Summarily, reduced dimensionality only occurred during NREM SWRs in the R2A circuit. All other conditions either exhibited no change or an increase in dimensionality.

This is consistent with our hypothesis that the RSPv, a DMN hub, exhibits state-/circuit-dependent vulnerability to APP/A*β* such that R2A is degraded while R2V is not. However, there is an simple alternate explanation: our results simply illustrate a failure in dACC. To address this, we took advantage of the directionality of our analyses. We switched each source and target, allowing us to examine a second set of circuits: V2R (VISp to RSPv) and A2R (dACC to RSPv) (Fig. 5A). In contrast to the results presented above, A2R dimensionality exhibited no differences between WT and APP dimensionality in any condition (NREM SWRs: p = 0.206, delta NREM: p = 0.644, waking SWRs p = 0.166) (Fig. 5B-D, right). Nor were there were significant differences between WT and APP animals in V2R dimensionality in any condition (NREM SWRs: p = 0.581, delta NREM: p = 0.287, waking SWRs p = 0.141) (Fig. 5B-D, left). Apparent differences in the raw distributions (inset histograms) were driven by individual animals, an effect captured and accounted for by linear mixed model (Figure S9). These data suggest that deficits are driven by a computational subspace of RSPv, and not explained by a failure of dynamics in ACAd.

**Figure 5:**
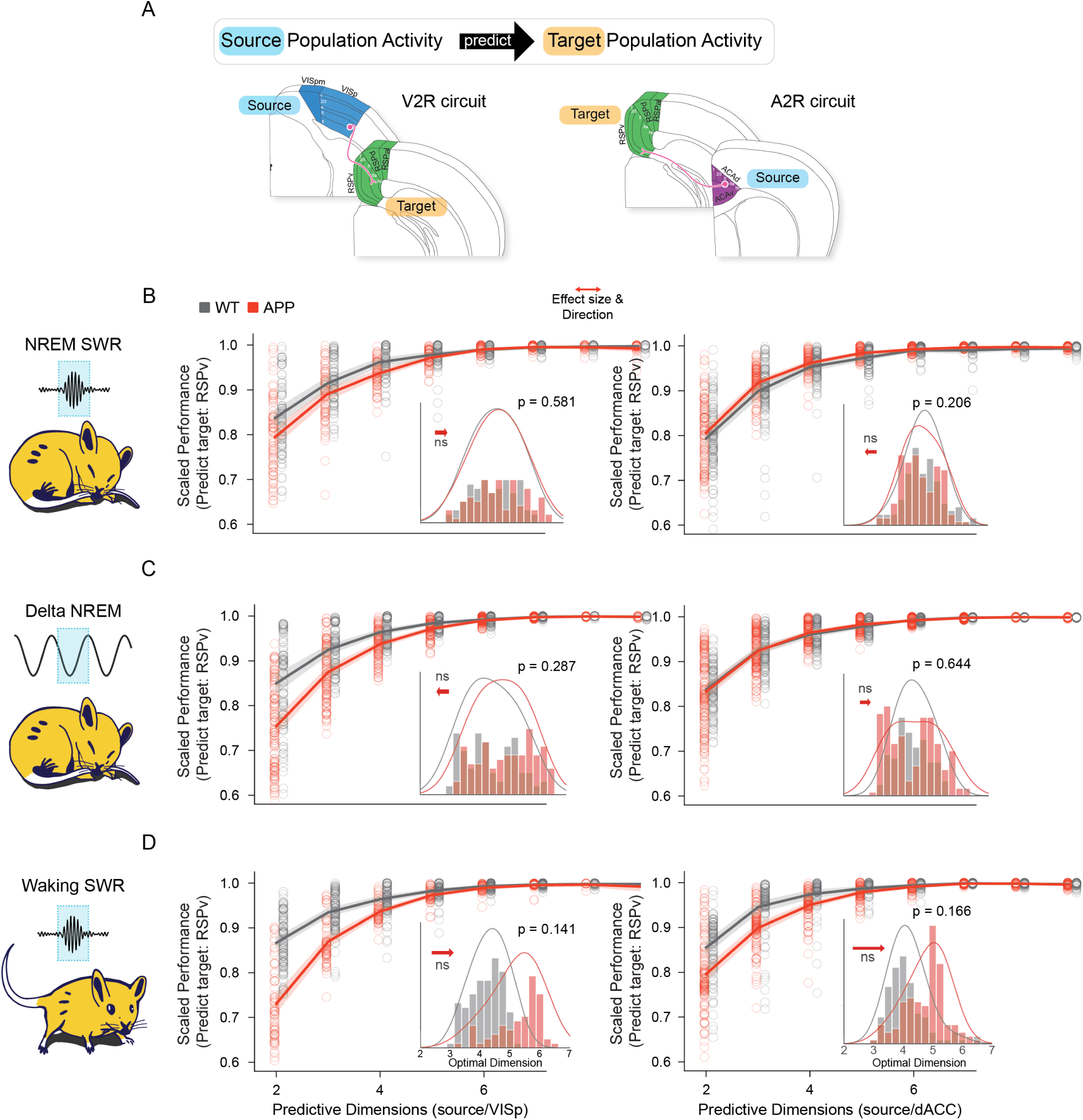
Switching source and target reveals no significant changes in communication subspace dimensionality between APP/PS1 and WT mice. Data are analyzed as in Figure 4, but the regions used for source and target were reversed to generate two new circuits: V2R and A2R. Dimensionality of communication subspace in these circuits was measured. **A.** (top) Conceptual diagram of analysis. The activity of an ensemble of single neurons in a source region is used to predict the activity of neurons in the target region. The dimensionality of the source ensemble is then progressively reduced, and the loss of predictive power is measured. (bottom) Diagram of the A2R circuit and V2R. In each case, the target is the same: the RSPv. **B.** Dimensionality of V2R and A2R circuits in NREM sharp wave ripples (SWRs) as a function of genotype. Cartoon illustrates the behavioral/neurophysiological state. The left data column is from V2R, the right column is A2R. As the dimensionality of the source data is experimentally reduced, the predictive performance (scaled to proportion of the full prediction) is plotted. Each point represents one bootstrap, and the fitted line is a mixed model regression fit (logit(scaled performance) genotype * circuit * state + (animal/date/epoch/bootstrap). The last term represents nested random effects, see methods). The inset shows distribution of the optimal dimension (defined as the number of dimensions required to explain 90% of the full interaction) by genotype. The red arrow shows the direction of the effect (WT-APP) and the effect size (length of arrow). **C.** Same as B but d_1_u_3_ring randomly identified, slow wave (delta: 0.1-4 Hz) enriched, non-SWR intervals of NREM sleep. **D.** Same as B but in SWRs occurring during waking.

Across four circuits, the only incidence of reduced dimensionality occurred during NREM SWRs in the R2A circuit. This demonstrates that the functional consequence of APP/A*β*-related vulnerability may not be strictly reducible to structural and anatomical degeneration. Structural/anatomical changes do not vary on the timescale of hundreds of ms (e.g., SWRs). Taken in the context of global A*β*, global APP overexpression, and global presenilin overexpression, these data suggest that demonstrate that selective vulnerability is not explained by proximity to disease-related molecules. Computationally-defined subsets of key circuits are disrupted in a dynamic and state-dependent fashion, consonant with substrates related to fast patterning and the routing of information, such as neuromodulatory systems and recurrent inhibition^43,44^.

## DISCUSSION

The brain regions comprising the default mode network (DMN) are selectively vulnerable to A*β* deposition and Alzheimer’s Disease (AD) -related disruption. This vulnerability has been widely described, particularly in human clinical populations, yet it has been challenging to separate two possible explanations of this phenomenon. The simplest case is that the pattern of A*β* deposition early in AD determines the pattern of affected circuits. In other words, A*β* plaques are toxic to nearby neurons, and plaque expression is limited to the DMN. Alternatively, early AD could represent the confluence of two events. Specific circuits are intrinsically vulnerable to A*β*-related damage, and it is in these circuits that A*β* takes hold. To directly test which of these explanations is more likely, we made four-site recordings of neuronal ensembles in individual APP/PS1 mice, which are characterized by a widespread A*β* burden across the isocortex and hippocampus. We recorded two key circuits, one previously defined as in-DMN and one previously defined as out-DMN^10^. Each of these shares the same source (RSPv, a DMN hub) but differs in their targets (dACC, a DMN region, and VISp, a non-DMN region). Our data reveal circuit level-differences in the impact of A*β* (and global APP/PS1 overexpression), despite both widespread A*β* load (and pan-neuronal APP/PS1 overexpression) and the fact that the same ensembles comprised the source of each circuit. Not only do our data offer a unique resolution of intraregional circuitry in an animal model of amyloid overexpression (a key feature of AD), but our recordings span 24 h of unrestrained, natural behavior. In this context, disruptions of the R2A (in-DMN) circuit were exaggerated throughout NREM sleep, and showed robust effects during sharp wave ripples (SWRs), a neurophysiological event with consequences for memory consolidation.

While high-density, multi-region recordings of single units in freely behaving animals offer powerful insights into complex neuronal interactions during behaviors, there is a tradeoff between resolution and the anatomical breadth of observation. Given the anatomical specificity of this approach, we carefully chose the specific in-DMN and out-DMN circuits, guided by prior work from our group^11^. Specifically, we sought a pair of in- and out-DMN circuits, each of which was monosynaptically connected to a common source. While our choice of circuits meets these criteria, it is currently impossible to simultaneously record single neuron activity with spike time resolution from all circuits originating in the DMN. As a result, our data offer support for the hypothesis that circuits originating and terminating in the DMN are intrinsically vulnerable, even in comparison to those with the same origins but out-DMN targets. However, extensive further work will be required to establish the generalizability of these findings. It is also important to acknowledge a caveat associated with the use of transgenic models of A*β* overexpression: it is possible that observed effects are driven by the transgene and not A*β* production and deposition. While prior work suggests that pathology in this context is mechanistically accounted for by A*β*^45,46^, our fundamental conclusions would not change if effects were related to transgene expression. Overexpression should be equally distributed in all circuits, thus still highlighting the intrinsic vulnerability of the in-DMN circuit.

Many mouse models of amyloid deposition, a key feature of AD, rely on the transgenic overexpression or knock-in of humanized APP carrying familial disease mutations that increase A*β* production. Lines differ in the anatomical distribution and timing of plaque deposition^11,47^, as well as the timing and extent of behavioral, functional, and cellular damage. The APP/PS1 mouse used in this study accumulates plaque in the cortex and hippocampus first with widespread coverage, yet minimal degenerative damage, at 12-16 months of age^12^. Similarly, we detected no significant differences in the yield of single units or histology as a function of genotype that might have indicated neuronal loss. The broad anatomical expression of A*β* in these mice is related to the promoter used to drive the transgenes; an important caveat that is often considered undesirable for modeling natural human disease progression. Further, it is worth noting that our conclusions wouldn’t change if APP/PS1 overexpression were responsible for the circuit-level disruptions observed here. Whether explained by APP/PS1 or A*β*, our data suggest that a subset of circuitry within the DMN is selectively/intrinsically vulnerable in the context of a global insult (whether that’s pan-neuron APP/PS1 over expression, or widespread tau deposition). Here we leverage this feature in a positive light to address an important question about circuit vulnerability.

While APP overexpressing mice exhibit relatively little neurodegeneration, there is evidence of neuronal dysfunction by 12 months in the APP/PS1 line. These animals experience frequent spontaneous epileptiform discharges on EEG as well as seizures, which we noted in our observations of behavior and physiology^48^. In the APP/PS1 mouse, A*β* has been shown to increase neuronal activity^23,24^ as well as decrease neuron firing, perhaps explained by a neuron’s proximity to nearby plaques^25^. This may also depend on brain state or context^49,50^, highlighting the need to examine neuronal activity in multiple regions across a range of naturally occurring conditions. At a lower spatial resolution, APP overexpression reduces the power of gamma oscillations^51^ and alters coupling between the hippocampus and cortex during sleep^37^. Our results suggest that alterations in activity are likely to be circuit-specific and may require simultaneous monitoring of multiple populations of neurons.

The major impact of pathology, especially early in disease progression, may vary throughout the circadian cycle and depend on changes in brain state and behavior. Our data emphasize a key role for brain state in understanding amyloidosis. Specifically, our results reveal that, in the in-DMN circuit, dysfunction is pronounced during NREM sleep. This is of particular interest in light of the relationship of NREM sleep to memory. Much work has linked sleep, particularly NREM, to memory consolidation^52^ often involving hippocampal-cortical interactions^53^. Perhaps it is unsurprising then that early AD in humans is marked by disrupted sleep structure^54–56^. This is likely a bidirectional interaction, as chronically disrupted sleep is predictive of disease^57^. Similarly, APP/PS1 mice show disrupted sleep/wake cycles^58^. Even within sleep, pathology may be best understood in the context of discrete events. Our data suggest that, in APP/PS1 animals, the timing of SWR driven activity is disrupted in cortical regions that comprise the DMN but is intact in non-DMN cortices. In the context of NREM sleep, peak firing rates during SWRs occurred earlier in RSPv and later in ACAd in APP mice compared to WT. This suggests that the impact of APP/A*β* on higher order features of neuronal activity, such as timing and patterning, is not a trivial effect of, for example, hyperexcitability^48^. Subtle disruptions of the dynamics of specific transcriptomic cell types^59^ and/or variable expression of compensatory plasticity^16,20,60–62^ are plausible examples by which A*β* could drive complex and circuit-specific alterations in neuronal dynamics.

Notably, disruption of SWRs in the R2A circuit, both in terms of peak timing and dimensionality, is specific to state; SWRs that occur during quiet waking appear to be largely unaffected by APP/A*β*. This is less surprising than it may seem at first glance. SWRs in wake and NREM have distinct neuromodulatory contexts and cognitive implications: waking SWRs are associated with planning future actions and memory retrieval, whereas NREM SWRs are linked with memory consolidation^63^.

Further, the dimensionality of communication in the in-DMN circuit is significantly reduced during NREM SWRs while the out-DMN circuit is equivalent to WT. The suggestion that SWRs are a locus of pathological effect is consistent with observations of abnormal SWR properties in mouse models of a range of neurodegenerative diseases^37,64,65^. In the APP/PS1 RSPv, our results reveal an interesting detail: neuronal firing is elevated during SWRs but not during general NREM sleep. This suggests that RSPv is not tonically hyperactive but has exaggerated responses when activated, consistent with impaired feedforward inhibitory input from CA1^39,40^. Further work is required to test the hypothesis that disruption in the balance of excitation and inhibition^51^ may be restricted to brain states or events^66^.

Our findings align well with the cascading network failure hypothesis in AD^67^. This asserts that AD pathogenesis is the result of network failures, beginning in the posterior DMN, that shift processing burden to the next step of the system, which subsequently fails as the damage propagates along lines of high connectivity (i.e. through the DMN, initially). This model predicts increases in connectivity that precede structural and functional disruption, and accounts for the prion-like spread of A*β*^55,68^. Key to this theory is that altered functional communication between DMN regions is an inciting event. Consistent with this, our data confirm impaired communication in the in-DMN pathway, despite equivalent amyloid burden in the off-DMN pathway. Also of note, our data reveal lagged connectivity in the direction of RSPv to ACAd (posterior to anterior DMN), which is also consistent with data from humans with AD^67^. The consistency of our observations and key features of human pathology suggests that the mechanisms underlying the selective vulnerability of DMN circuits are phylogenetically conserved and are not tethered to the specifics of the disease process.

We propose that in AD, the selective vulnerability of the DMN is a consequence of intrinsic vulnerability to injury and is not explained by the anatomical specificity of amyloid accumulation. Within DMN regions, the vulnerability of individual neurons to functional disruption is likely to be determined by the cell type specific gene expression underlying connectivity patterns^10,69^. How this interacts with the increased activity in the DMN at rest is a crucial question, as neuronal activity can exacerbate A*β* pathology^70^. The advent of new technologies to measure mesoscale connectivity, record neuronal activity, and resolve spatial transcriptomics will make it possible to further disentangle cellular and circuit rules by which disease spreads through the DMN.

**Figure S1:**
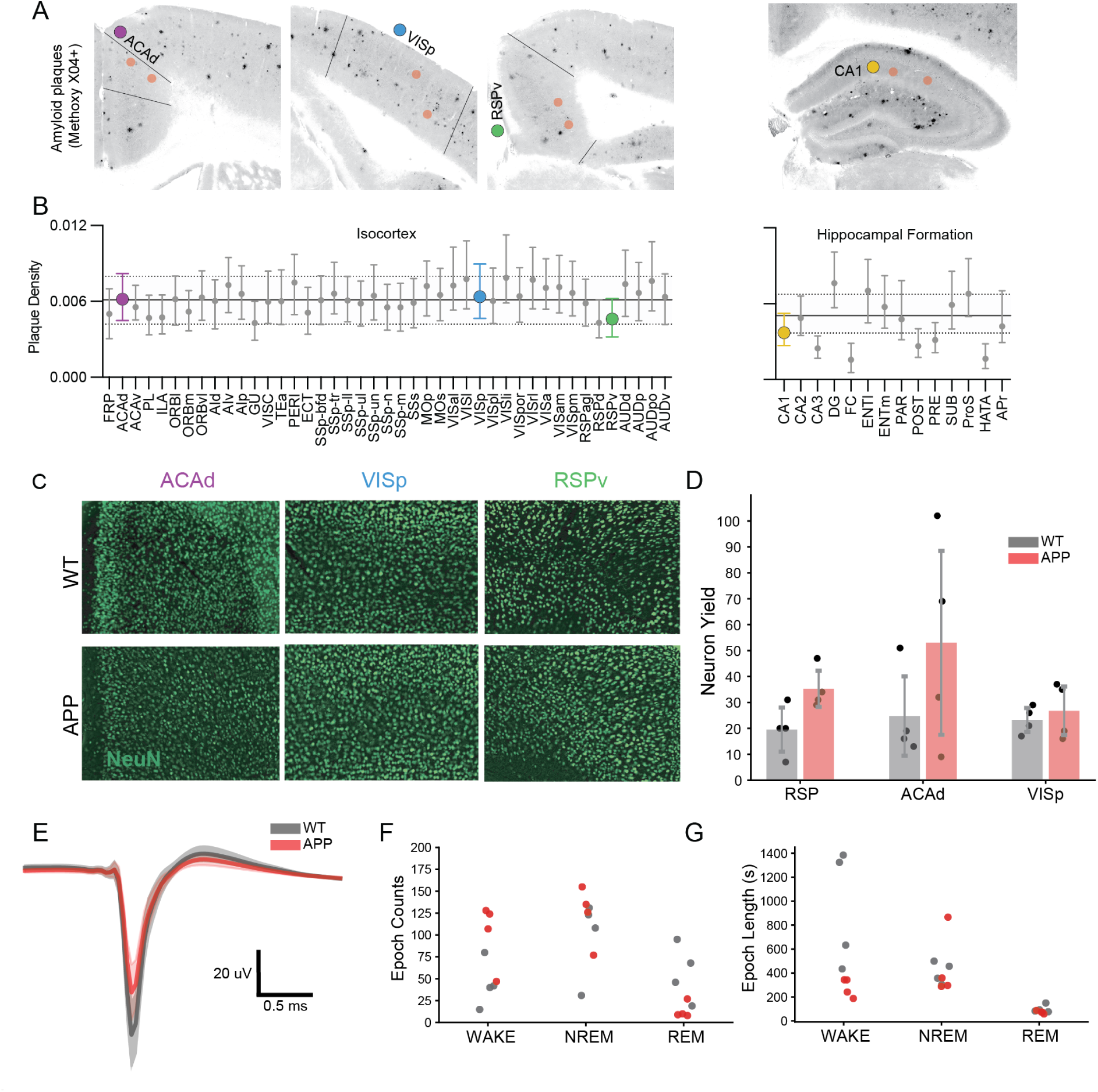
Quantification of amyloid isocortical and hippocampal plaque burden and recording context. Plaque density across all cortical and hippocampal brain regions of 12–15-month-old APP/PS1 mice. Plaques were labeled with methoxy-X04 and imaged across the entire brain using serial two-photon tomography. Plaque density is calculated as the fraction of brain structure volume containing plaques. **A.** Representative images reveal labeled plaques in the three isocortical and hippocampal sites where electrodes were implanted in this study. Red circles indicate reconstructed electrode implantation sites from two animals. **B.** Plaque densities measured with automated segmentation and registration for n=160 12–16-month-old APP/PS1 mice are plotted for all 43 isocortical and 14 hippocampal formation regions annotated in the Allen CCFv3^11^. Plots show the median and interquartile range (IQR). Solid and dashed lines show the median plaque density and IQR for the entire isocortex or hippocampal formation. **C.** Representative NeuN histology from each genotype. **D.** Single unit yield for each recorded region as a function of genotype. No significant differences were found, although RSPv trended to yield more neurons in APP/PS1 animals than WT (n = animal; Wilcoxon rank sum p = 0.061 RSP, p = 0.386 ACAd, p = 0.773 VISp). **E.** Average spike waveform for regular spiking units (presumptive excitatory neurons). Color indicates genotype. **F,G.** The number and mean duration of epochs of wake, NREM, and REM per animal, color indicates genotype.

**Figure S2:**
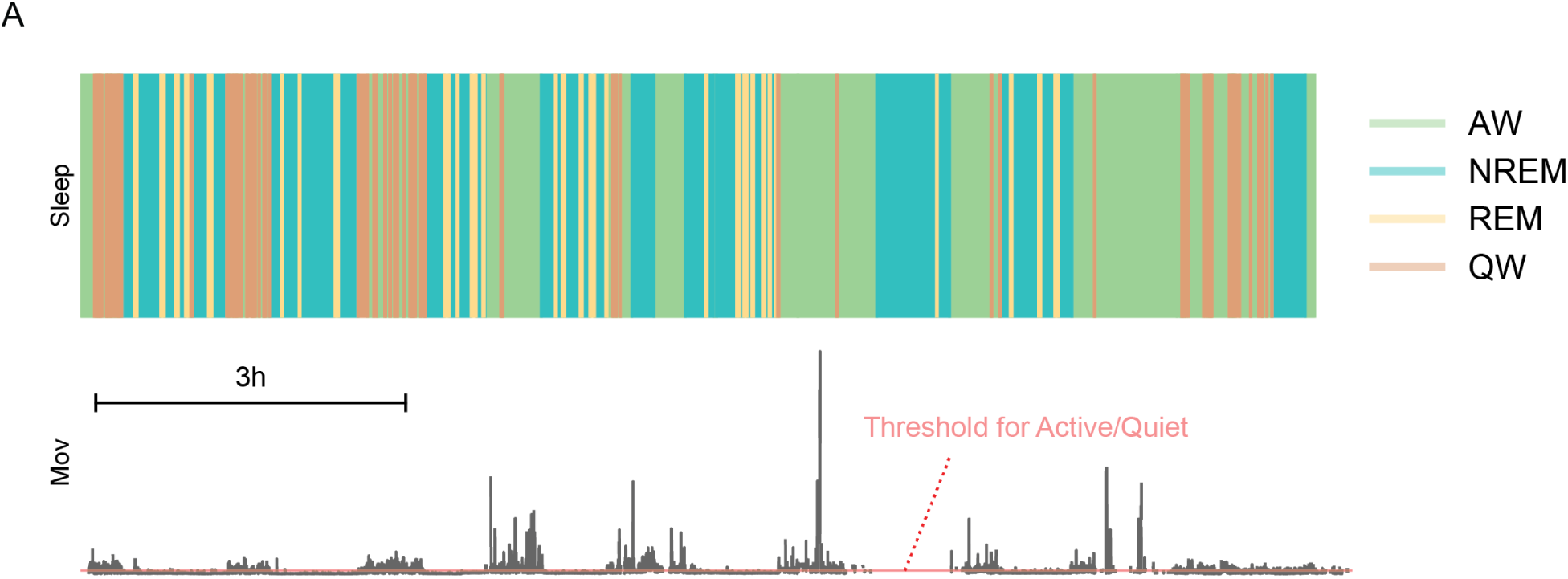
Thresholding of movement to determine active wake and quiescence. **A.** Area plot of 12 h of sleep/wake states scored by a human expert (see methods). Periods of waking reflect subsequent classification as active (AW) or quiescent (QW). **B.** Raw movement trace derived from the output of a convolutional neural network trained to track animal pose in individual video frames. Dashed red line indicates the threshold used to delineate periods of quiescence from those of activity. The threshold was established empirically for each animal to account for differences in camera angles and individual behavioral patterns. Activity assessments during waking epochs were manually confirmed to eliminate occasional model error.

**Figure S3:**
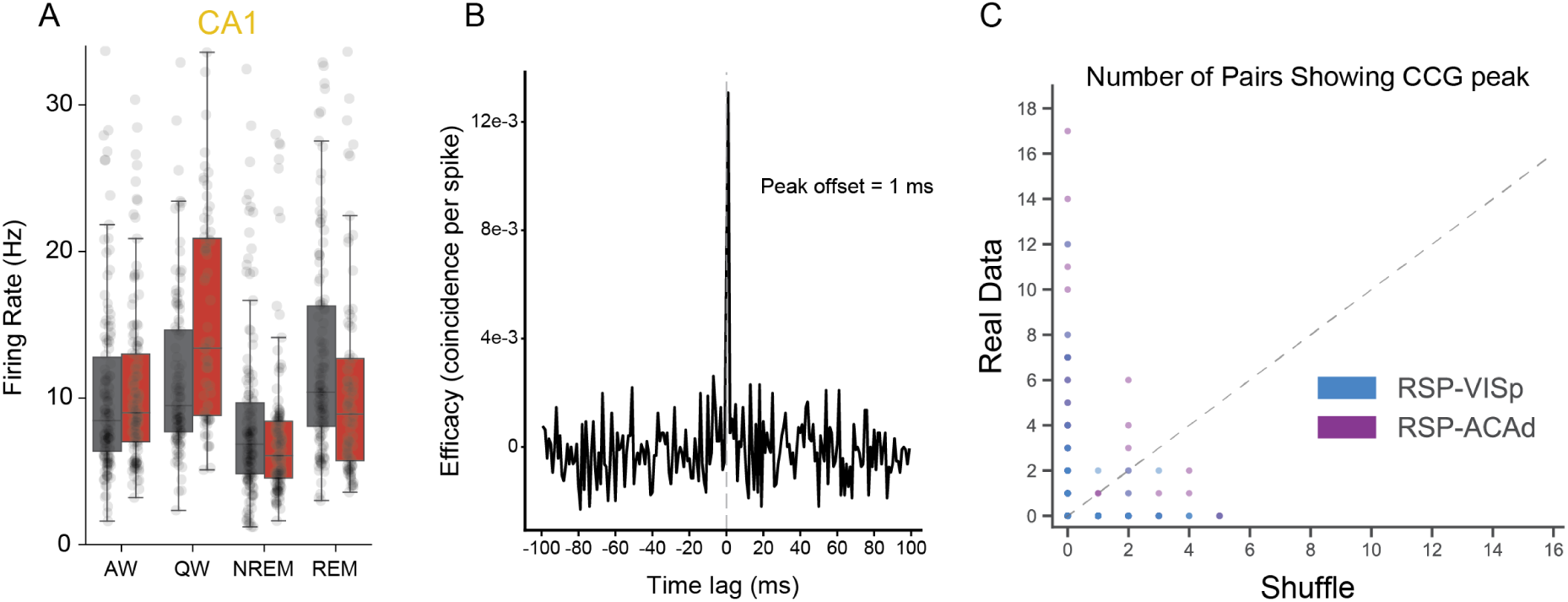
Single unit firing rate in CA1 and electrophysiological evidence of monosynaptically connected pairs connecting RSPv to both VISp and ACAd. **A.** Mean firing rate of single units in CA1 as a function of genotype and brain state. No significant differences were observed (AW p = 0.46, QW p = 0.7, NREM p = 0.169, REM p = 0.119, linear mixed effects regression with animal and recording epoch as nested random effects). **B.** Example cross correlation of spike timing between a pair of single units, one in RSPv and the other in ACAd. The sharp peak in correlation at 2 ms is consistent with a monosynaptic delay. **C.** Quantification of the number of single unit pairs whose correlations are consistent with monosynaptic connectivity. Pairs either span RSPv and VISp (blue) or RSPv and ACAd (purple). Each axis denotes the number or pairs identified with a time lag of less than 10 ms. Each plotted point represents the number of monosynaptic pairs identified in one dataset. Intact pairs (Y axis) are plotted against shuffled data (X axis) to control for the probability of observing low-latency correlations due to chance in population recordings. The majority of datasets revealed no pairs in shuffle and 1-18 pairs in the intact data (left most column)

**Figure S4:**
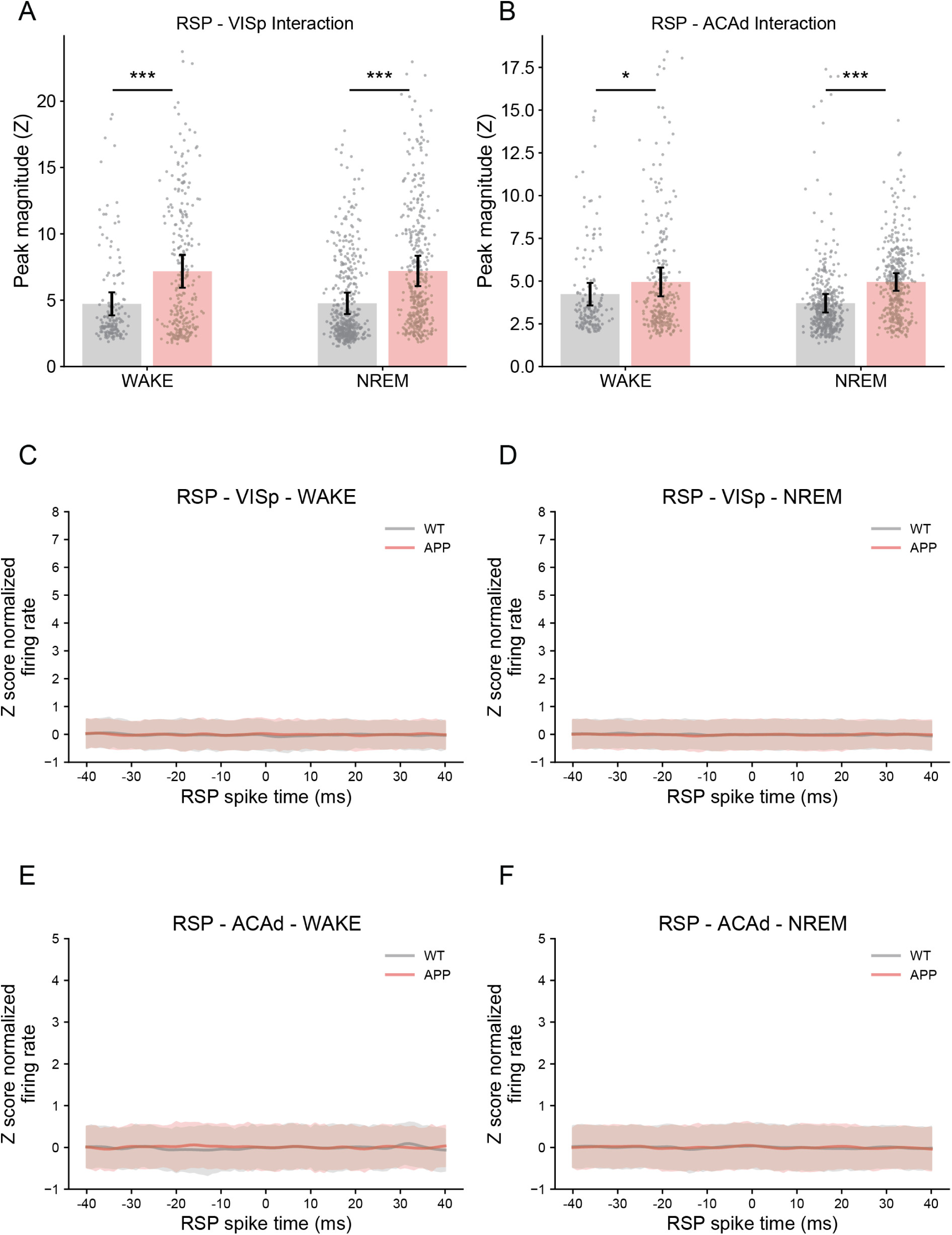
Magnitude of correlation peak by circuit and shuffle control. **A.** Peak magnitude of the correlation of z normalized neuronal firing rates in wake and NREM (n = epochs) for RSPv-to-VISp lagged correlation (Wilcoxon rank sum with Bonferroni correction p = 3e-06 Wake, p = 0.00 NREM). **B.** Peak z normalized correlation in wake and NREM (n = sleep/wake epochs) for RSPv-to-ACAd lagged correlation (Wilcoxon rank sum with Bonferroni correction p = 0.018 Wake, p = 0.00 NREM). **C-F.** Lagged correlation between source RSPv activity and shuffled target data. The target is indicated in the subplot title, as is the brain state.

**Figure S5:**
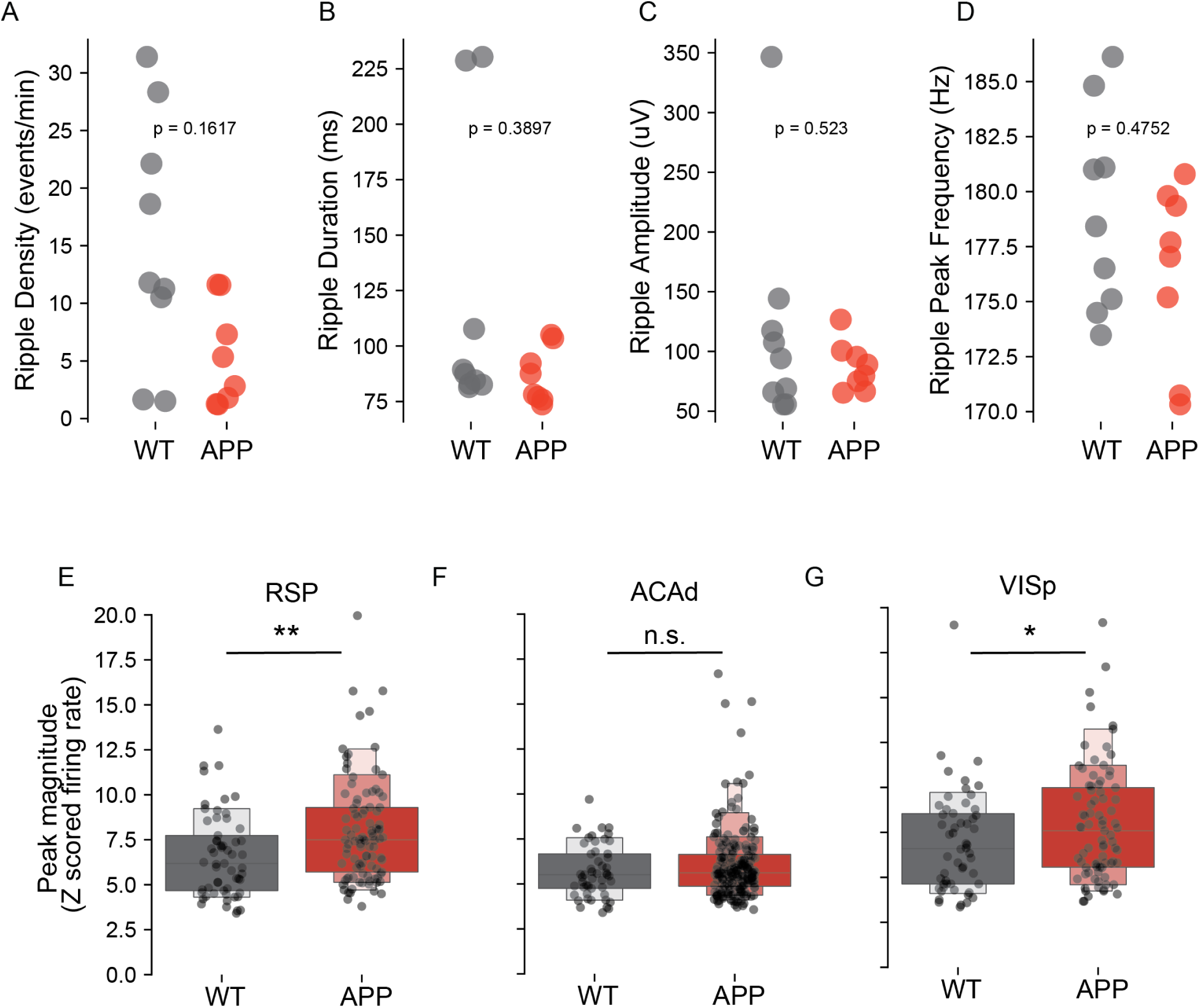
Sharp wave ripple statistics as a function of genotype. **A-D.** Basic features of sharp wave ripples in WT (gray) and APP/PS1 (red) animals. No significant differences were observed in ripple density (A), duration (B), amplitude (C), or peak frequency (D), linear mixed effects model: ripple feature genotype + (animal), where the last term represents a random effect of individual. n = recording epoch. **E-G.** Peak magnitude of z scored firing rate around sharp wave ripples in RSPv (E), ACAd (F), and VISp (G). (n = single units, * p *<* 0.05, ** p *<* 0.01, n.s. not significant. Wilcoxon rank sum).

**Figure S6:**
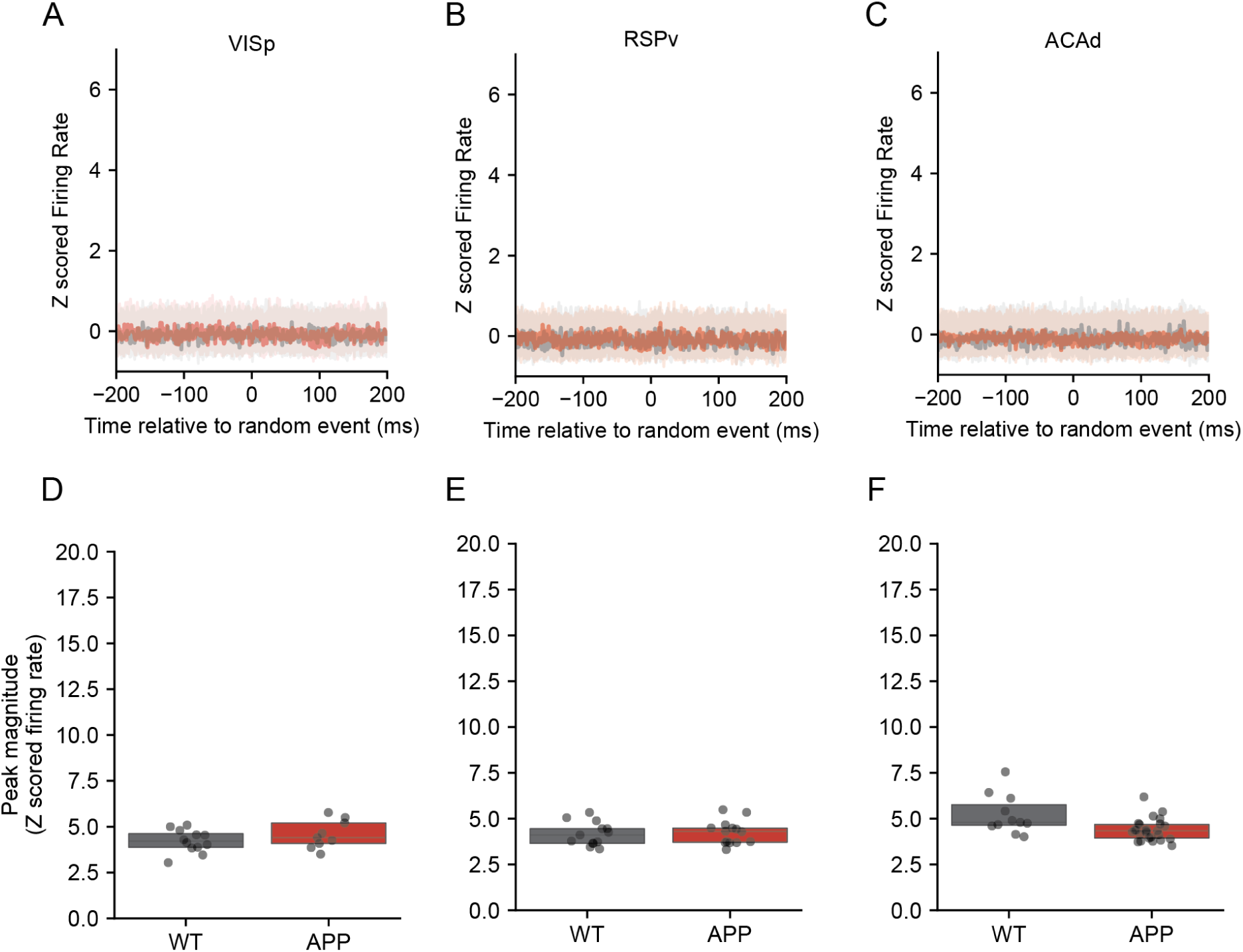
Peri-event activity outside of sharp wave ripples. To understand the specificity of the The sharp wave ripple associated activity in VISp, RSPv, and ACAd is absent outside of sharp wave ripples. **A-C.** Z scored activity aligned to random event times (outside sharp wave ripples) in A VISp, B RSPv, and C ACAd. **D-F.** Peak magnitude of z scored firing rate around random event times (non-sharp wave ripple) (n = single units, * p *<* 0.05, ** p *<* 0.01, n.s. not significant Wilcoxon rank sum) in D VISp, E RSPv, and F ACAd.

**Figure S7:**
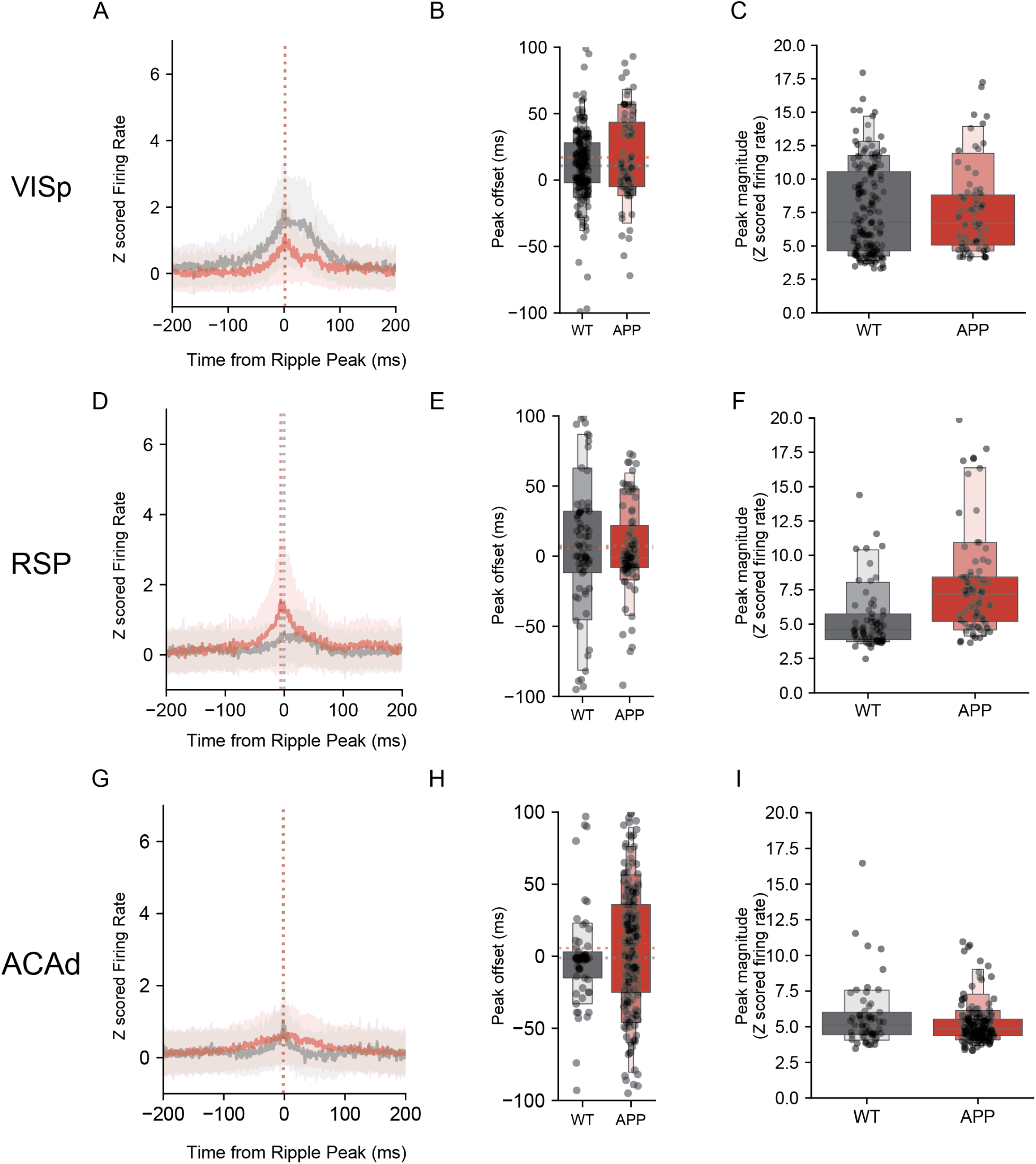
Peri-sharp wave ripple activity during wake by genotype. Average single unit responses to waking sharp wave ripples in **A** VISp, **D** RSPv, and **G** ACAd. The timing of single unit activity during wake SWRs does not differ by genotype in **B** VISp, **E** RSPv, or **H** ACAd. The magnitude of responses to wake SWRs in **C** VISp, **F** RSPv, and **I** ACAd.

**Figure S8:**
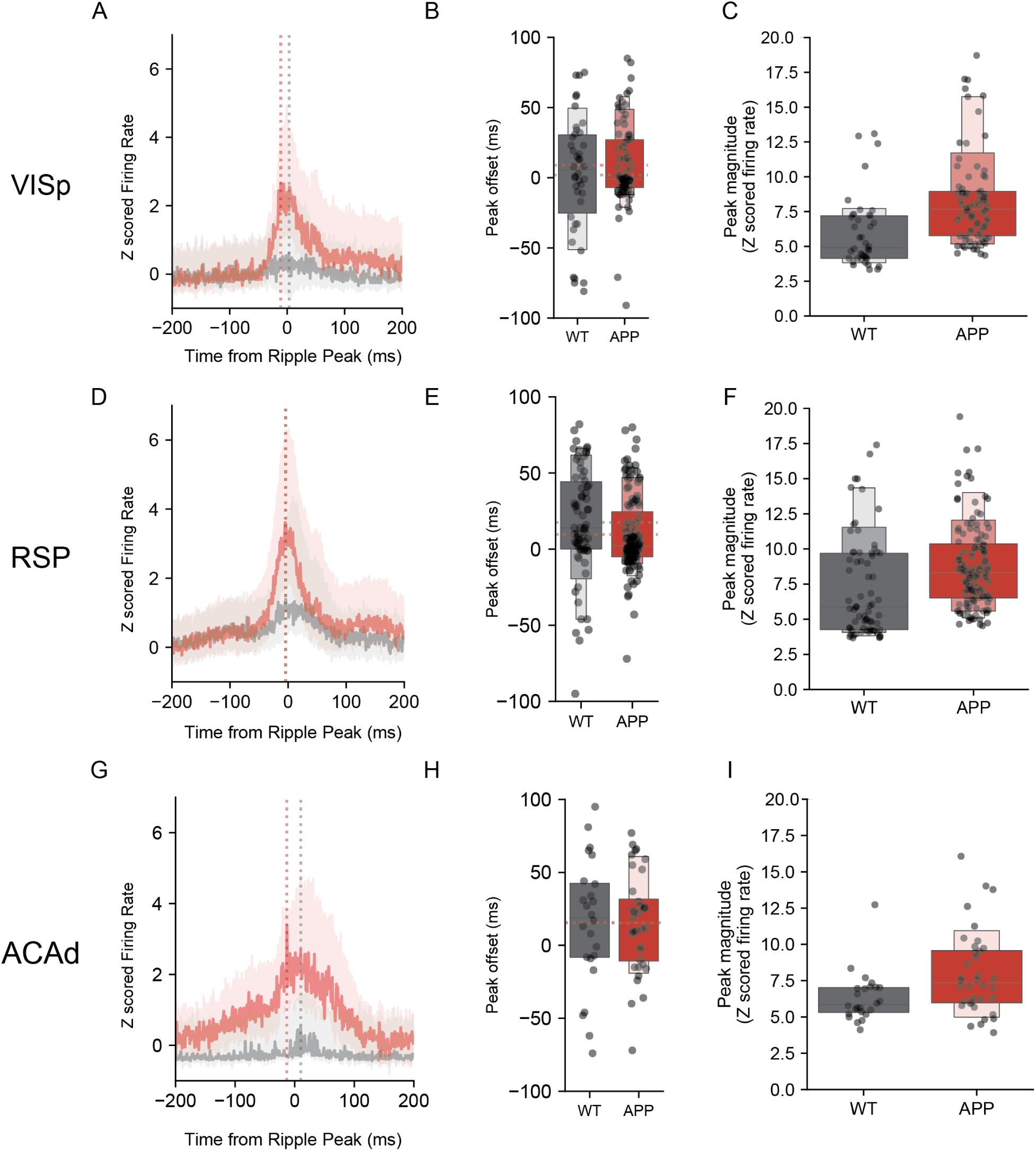
Peri-sharp wave ripple activity of FS cells during NREM. Average single unit responses of FS cells to NREM sharp wave ripples in **A** VISp, **D** RSPv, and **G** ACAd. The timing of single unit activity during wake SWRs does not differ by genotype in **B** VISp, **E** RSPv, or **H** ACAd. The magnitude of FS responses to NREM SWRs by genotype in **C** VISp, **F** RSPv, and **I** ACAd.

**Figure S9:**
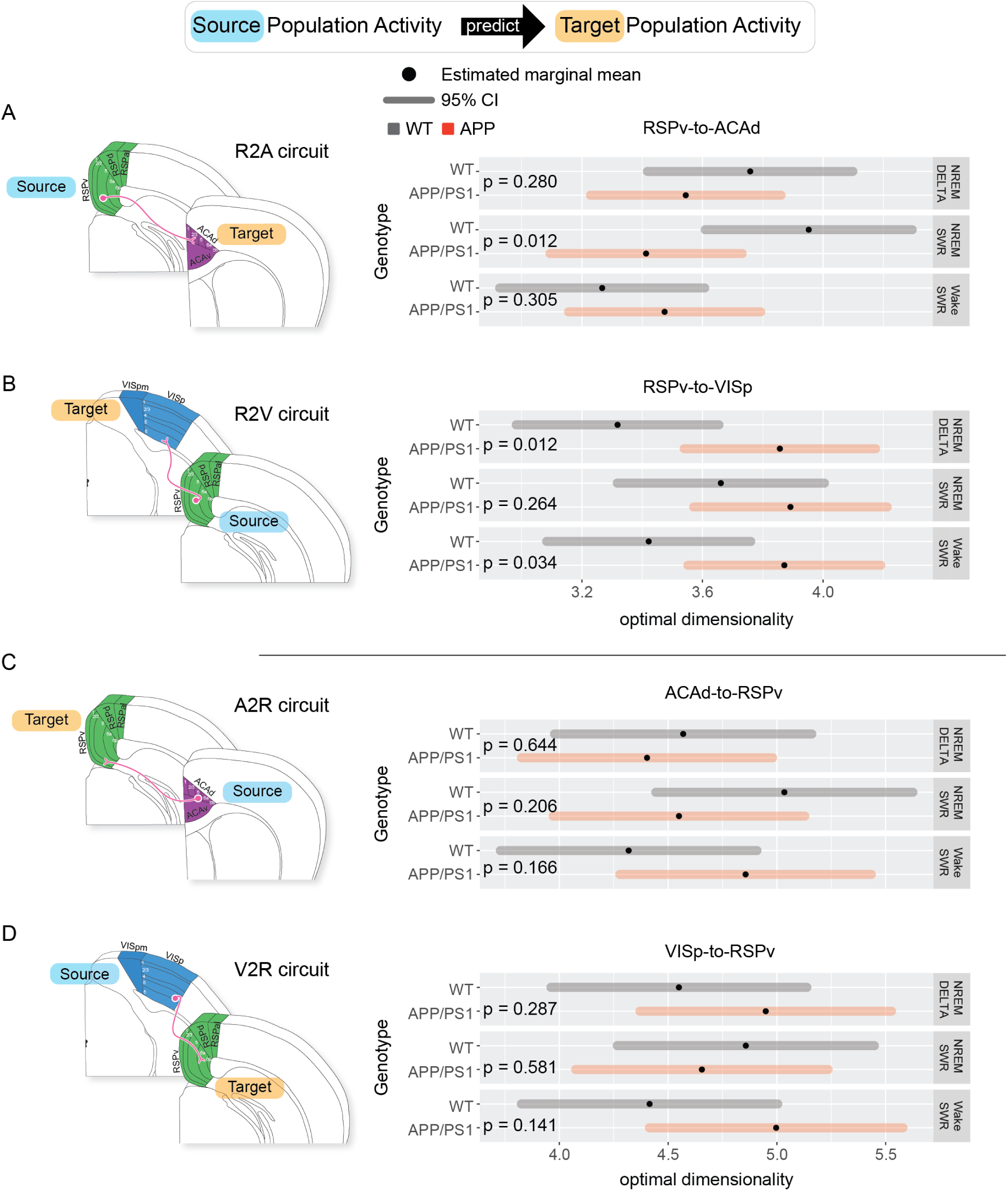
Estimated marginal means of communication subspace dimensionality in all four recorded circuits. (top) Conceptual diagram of analysis. The activity of an ensemble of single neurons in a source region is used to predict the activity of neurons in the target region. The dimensionality of the source ensemble is then progressively reduced, and the loss of predictive power is measured. **A** (left) Diagram of the R2A circuit. (right) Estimated marginal mean (black point) and 95% confidence interval (CI; shaded horizontal bar) of the linear mixed model by genotype (gray vs red) and state (row). Model: logit(scaled performance) genotype * circuit * state + (animal/date/epoch/bootstrap). The last term represents nested random effects, see methods). **B** Same as A, but for the R2V circuit. **C** Same as A, but for the A2R circuit. **D** Same as A, but for the V2R circuit.

## METHODS

### Animal husbandry and model

All procedures involving mice were performed in accordance with protocols approved by the Washington University in Saint Louis Institutional Animal Care and Use Committee, following guidelines described in the US National Institutes of Health Guide for the Care and Use of Laboratory Animals. We used 4 heterozygous APP+/− mice and 4 nontransgenic littermates (APP-/-) from the transgenic APP/PS1 line (B6.Cg-Tg[APPswe, PSEN1dE9]85Dbo /Mmjax, MMRRC Stock No: 034832-JAX^71^). All mice were between 12-16 months of age at time of experiment. Mice were housed in an enriched environment and kept on a 12:12 h light:dark cycle with ad libitum access to food and water.

### Surgical procedures

All mice underwent multisite electrode array implantation surgery. Mice were anesthetized with inhaled isoflurane (1-2% in air) and administered slow release buprenorphine (ZooPharm, 0.1 mg kg-1) for analgesia. The mouse’s skull was secured in a robotic stereotaxic instrument (NeuroStar, Tubingen, Germany), and the skin and periosteum covering the dorsal surface of the skull was removed. Lambda and bregma midline positions were identified and used as reference coordinates for alignment with stereotaxic atlases using Neurostar StereoDrive software. Four craniotomies (diameter 1-1.5 mm) were drilled over target implantation sites using the automatic drilling function of the stereotaxic robot, and the dura mater membrane was resected. Four brain regions were implanted with custom 64-channel nichrome tetrode-based arrays cut at a 45*^◦^* angle such that the distance between the shortest and longest tetrode was 350-500 *µm*. Arrays were fixed (not drivable) and separated from headstage hardware by a flex cable, thus allowing an arbitrary geometry of multiple probes. In each mouse, four 64 channel arrays were targeted so that the longest recording wire was centered on the following stereotaxic coordinates (AP/ML/DV relative to bregma and dura, in mm): CA1 (−2.52/−1.75/1.5), ACAd (0.6/−0.25/1.15), RSPv (−3.28/− 0.95/1.17), and VISp (−4.03/−2.88/1.25). Electrode bundles were lowered into brain tissue at a rate of 3 mm/min using a custom built vacuum holder. Anatomical location was confirmed post hoc via histological reconstruction. Arrays were secured with dental cement (C&B-Metabond Quick! Luting Cement, Parkell Products Inc; Flow-It ALC Flowable Dental Composite, Pentron), and headstage electronics (eCube, White Matter LLC) were bundled and secured in custom 3D-printed housing. Four-module (256 channel) implants weighed approximately 2 g. Post surgery, mice were administered meloxicam (Pivetal, 5 mg kg-1 day-1 for three days) and dexamethasone (0.5 mg kg^−1^ day^−1^ for three days) and allowed to recover in the recording chamber prior to recording.

### Recording procedures

Recordings were made using custom tetrode-based arrays. Sixteen tetrodes (64 channels) were soldered to a custom-designed PCB (5 mm x 5 mm x 200 *µm*) which stacked horizontally with a similarly sized amplifier chip (White Matter LLC, Seattle). PCB/amplifier pairs further stacked with 3 additional pairs (total 4 modules, 256 channels). Recordings were conducted in an enriched home cage environment with social access to a litter mate through a perforated acrylic divider. Freely behaving mice were attached to a custom built cable with in-line commutation. Neuronal signals were amplified, digitized, and sampled at 25 kHz alongside synchronized 15-30 fps video using the eCube Server electrophysiology system (White Matter LLC). Recordings were conducted continuously for between two weeks and three months. Data and video were continuously monitored using Open Ephys^72^ and Watch Tower (White Matter LLC). 24 h blocks of data were identified for inclusion in these studies first by the absence of hardware and/or software problems, for example cable disconnects or dropped video frames, respectively, and second by the maximal yield of active channels. Beyond these criteria, selection of 24 h blocks was arbitrary.

For experiments involving spike-sorted data, raw data were bandpass filtered between 350 and 7,500 Hz and spike waveforms were extracted and clustered using a modified version of SpikeInterface^73^ and MountainSort4^14^ with curation turned off. A custom XGBoosted decision tree was used to identify those clusters constituting single units. Clusters identified as single units were manually inspected to confirm the presence of high amplitude spiking, stable spike amplitude over time, consistent waveform shapes, and little to no refractory period contamination. For all downstream analyses, we excluded neurons with firing rates *<* 0.5 Hz or presence ratio *<* 0.8 throughout the duration of a sleep/wake epoch. RSU and FS units were classified using trough-to-peak times of spike waveforms: units with a trough-to-peak time of less than or greater than 0.4 ms were classified as FS or RSU units, respectively.

### Probe Localization

Following recording, mice were perfused with 4% formaldehyde (PFA), the brain was extracted and immersion fixed for 24 h at 4*^◦^* C in PFA. Brains were then transferred to a 30% sucrose solution in PBS and stored at 4*^◦^* C until brains sank. Brains were then sectioned at 50 *µm* on a cryostat. Sections were rinsed in PBS prior to mounting on charged slides (SuperFrost Plus, Fisher) and stained with cresyl violet. Stained sections were aligned with the Allen Institute Mouse Brain Atlas (Allen Institute for Brain Science, 2011) and tetrode tracks were identified under a microscope (Fig S1A).

### Sleep wake scoring

LFP was extracted in the 0.1-60 Hz range from 5 manually selected channels for each subject. Channels were averaged together and then spectrograms were generated in 1-hour increments. Sleep scoring was performed semi-manually in custom GUI software based on spectrogram data and movement tracking of the headstage generated with DeepLabCut^19^. States were assigned to NREM, REM, or Wake (Quiet, Active) based on delta bandpower (high in NREM), theta bandpower (high in REM), and movement/broadband LFP activity (high in wake) using well established methods^59,74^. A random forest trained on sleep scores from multiple human raters was used to assign sleep scores in 4 s epochs from spectrogram and movement data. These were checked manually by human raters and corrected where appropriate. For periods where sleep state could not be easily discerned by spectrogram and movement alone, temporally aligned video was consulted. For downstream analyses, we only included sleep/wake epochs of duration longer than 20 s to avoid epochs dominated by the effects of state transitions.

### Single Unit FR and Correlation Analysis

For each single unit, the average number of spikes was calculated in 1 s bins across each epoch. Then the mean firing rate was calculated across all epochs of a particular sleep state for each neuron individually. For each sleep epoch, spikes were binned in 1 ms windows for source and target regions. The Pearson correlation was calculated between the binned spike time series for each source-target pair of units at lags from −80 to +80 ms to generate a lagged correlogram. The mean of all lagged correlograms for every pair of source-target units was calculated for each epoch. Only epochs of greater than 60 s duration were included, after which correlograms were z-scored from the mean and standard deviation of the 20 ms flanks of the correlogram (−80 : −60 ms, +60 : +80 ms). Z-scored correlograms were then smoothed with 2 ms Gaussian kernel for visualization. For each epoch’s lagged correlogram, the peak position was found by identifying the time lag at which the maximum of the correlogram trace occurred on the z-scored correlogram in a 40 ms window around the center (−20 : +20 ms).

### Ripple Detection

For ripple detection, data from CA1 probes was downsampled to 1500 Hz and notch filtered at harmonics of 60 Hz to eliminate 60 cycle noise (60, 120, 180, 240, 300 Hz)^75^. The four channels with the lowest contamination in the ripple band were selected by determining the lowest mean bandpower in the ripple band (150-250 Hz) in 5 minute windows. Signal from the four selected channels was then bandpass filtered between 150 and 250 Hz before performing ripple detection with a publicly available Python package github.com/Eden-Kramer-Lab/ripple_detection implementing ripple detection from Karlsson and Frank^36^. Ripple amplitude was calculated as the difference between the maximum and minimum values of the ripple band-filtered LFP signal during each ripple. Only ripples with amplitude *>* 50 *µV* were included in downstream analyses. Ripples less than 100 ms apart were combined.

To calculate peri-event time histograms (PETHs) around sharp-wave ripples, single units were separated by putative cell type using spike waveform characteristics (RSU and FS). The PETH was then calculated by summing all spike events for each unit in a given region in a 1 s bin around each ripple time occurring during a NREM epoch. PETHs were then averaged across ripples for each unit. Average PETHs were then z scored using the flanks of the PETH (−30% : +30%) to enable comparison across animals/recording epochs. All z scored PETHs from all single units of a given cell type and region were then averaged together. To ensure that only responsive cells were included in genotype comparisons, only PETHs with activity two standard deviations above PETH flank activity were used in statistical comparison.

### Communication Subspace Analysis

For each recording epoch and animal, spike data from each brain region was aligned around instances of each of three neurophysiological phenomena: sharp wave ripples occurring during NREM, sharp wave ripples occurring during waking, and randomly selected intervals of NREM demonstrating slow waves (delta: 0.1-4 Hz) and lacking sharp wave ripples. For each type of event (NREM SWRs, waking SWRs, and delta NREM) spike times were binned (100 ms) in a 1 s window centered on each event (e.g., ripple). The average peri-event time histogram (PETH) across all events was subtracted from each individual PETH. Each PETH was then z scored to account for trial-to-trial variance, and then concatenated into one final time series for each recording epoch. In essence, we conducted standard, z-score normalized PETH analyses around SWRs or outside of SWRs, considering both NREM and wake.

From this preprocessed data, communication subspace analysis was performed using MATLAB code adapted from Semedo et al. 2019 (github.com/joao-semedo/communication-subspace)^42^. Briefly, reduced rank regression was performed between source RSP activity and target activity from VISp and ACAd. If data was rank deficient, non-contributory units were found via QR decomposition and removed. Only datasets with more than 10 units were used for subsequent reduced rank regression. To ensure that source and target contained equal numbers of units, units were randomly sampled from the larger dataset of either source or target data across 25 bootstraps to match the size of the smaller dataset. Reduced rank regression was then performed across 10 predictive dimensions using 10-fold cross validation. Optimal dimensionality was calculated as the number of predictive dimensions needed to perform within 1 standard deviation of performance at full rank^42^. Reduced rank regression was performed using the RidgeInit and Scaling parameters available in the Semedo MATLAB code. Normalized squared error was used for the loss function (sum of squared errors divided by the total sum of squares of the target data).

Results from each bootstrap were used in comparison of optimal dimensionality across genotypes. To ensure that analysis only contained reduced rank regression results where performance improved with increasing rank, RRR performance curves were included if performance at full rank was higher than performance at lowest rank and if performance at full rank was non-negative. The predictive dimensions – performance relationship curve was min-max scaled from 0 to 1 for each curve in order to make comparison across recording epochs and animals feasible.

### Statistical Analyses

Unless otherwise noted, all statistical analyses involved the implementation of a linear mixed-effects model using the lmer function from the lme4 package in R^26^. This model was developed to evaluate the potential influence of different factors on the relevant response variable (e.g., firing rate, predictive power). Fixed effects in models comprised main effects such as firing rate, genotype, circuit, and state, as well as their interaction effects. These variables represent distinct biological or experimental parameters in our study.

Random effects were also incorporated into the model to account for non-independence and potential clustering in the data. For example, we modeled random effects of animal, recordings, epochs, neuron ID, and bootstrap (where applicable) as random effects. When appropriate, we included nested random effects (such as bootstrap). These random effects account for the hierarchical or nested structure of the data and control for the non-independence of observations from the same animal, epoch, date, or bootstrap.

Model fitting was conducted using restricted maximum likelihood (REML) estimation, which provides unbiased estimates of variance and covariance parameters. After fitting the model, the fixed effect estimates were extracted to interpret the influence of each variable and their interaction effects on the response variable.

Data are reported as mean ± SEM unless otherwise noted. All tests were performed with a significance threshold set at p *<* 0.05.

## Data Availability

The datasets generated and/or analyzed in this study constitute ∼ 100 terabytes of raw neural (broadband) data. The data are stored in a cost efficient manner not immediately accessible to the internet. Data are available upon request.

## Code Availability

All relevant code from our lab, including software needed to run recordings like ours is in Python and is publicly available at github.com/hengenlab. Code necessary for the analyses used in this paper are available at github.com/hengenlab/appps1. Other groups’ code including Open Ephys, SpikeInterface, MountainSort4, DeepLabCut, RippleDetection, and Communication Subspace is publicly available as specified in Methods.

## Acknowledgments

This work is supported by NIH BRAIN Initiative R01NS118442 (KBH), 1R01EB029852 (ELD, KBH), BrightFocus Foundation Standard Award A2022038S (KBH), Cure Alzheimer’s Fund (KBH), Brain & Behavior Research Foundation Young Investigator Grant (29225 and in part by the National Institute on Aging R01AG047589 (J.A.H). We thank Erik Herzog, Charles Zorumski, and Erik Muziek for mentorship and consultation, Yifan Xu for assistance with sleep scoring pipeline, Gemechu Bekele and Anna Jasper (of the Adam Kohn Lab) for assistance with communication subspace analysis, and Ravi Chopra and Bryan Higashikubo for editing and review of the manuscript. We thank David Borchelt for sharing APP/PS1 mice.

## Author Contributions

S.B. assisted in recordings, performed data preprocessing, designed experiments, performed statistical analysis, and wrote the manuscript. J.N.M. conducted analyses, wrote the manuscript, and produced figures. C.F. performed surgeries and recordings. H.F. performed histology. K.B.N. provided technical consultation and software pipelines for data preprocessing. J.W. and J.A.H. contributed to initial concept, mentorship, and consultation. J.M.G. provided statistical assistance and helped construct figures. E.L.D. provided mentorship and assisted in writing the manuscript. K.B.H. lead and directed the project, produced figures, and wrote the manuscript.

## References

[1] Marcus E Raichle, et al. “A default mode of brain function.” In: Proceedings of the national academy of sciences 98.2 (2001), pp. 676–682.

[2] Heiko Braak, et al. “Amyotrophic lateral sclerosis—a model of corticofugal axonal spread.” In: Nature Reviews Neurology 9.12 (2013), pp. 708–714.

[3] Mathias Jucker and Lary C Walker. “Self-propagation of pathogenic protein aggregates in neurodegenerative diseases.” In: Nature 501.7465 (2013), pp. 45–51.

[4] Lary C Walker and Mathias Jucker. “Neurodegenerative diseases: expanding the prion concept.” In: Annual review of neuroscience 38.1 (2015), pp. 87–103.

[5] Yvette I Sheline, et al. “Amyloid plaques disrupt resting state default mode network connectivity in cognitively normal elderly.” In: Biological psychiatry 67.6 (2010), pp. 584–587.

[6] Federica Agosta, et al. “Resting state fMRI in Alzheimer’s disease: beyond the default mode network.” In: Neurobiology of aging 33.8 (2012), pp. 1564–1578.

[7] Andreas Horn, et al. “The structural–functional connectome and the default mode network of the human brain.” In: Neuroimage 102 (2014), pp. 142–151.

[8] Patric Hagmann, et al. “Mapping the structural core of human cerebral cortex.” In: PLoS biology 6.7 (2008), e159.

[9] Michael D Greicius, et al. “Resting-state functional connectivity reflects structural connectivity in the default mode network.” In: Cerebral cortex 19.1 (2009), pp. 72–78.

[10] Jennifer D Whitesell, et al. “Regional, layer, and cell-type-specific connectivity of the mouse default mode network.” In: Neuron 109.3 (2021), pp. 545–559.

[11] Jennifer D Whitesell et al. “Whole brain imaging reveals distinct spatial patterns of amyloid beta deposition in three mouse models of Alzheimer’s disease.” In: Journal of Comparative Neurology 527.13 (2019), pp. 2122–2145.

[12] Niels J Rupp, et al. “Early onset amyloid lesions lead to severe neuritic abnormalities and local, but not global neuron loss in APPPS1 transgenic mice.” In: Neurobiology of aging 32.12 (2011), 2324–e1.

[13] Fabrizio Trinchese et al. “Progressive age-related development of Alzheimer-like pathology in APP/PS1 mice.” In: Annals of Neurology: Official Journal of the American Neurological Association and the Child Neurology Society 55.6 (2004), pp. 801–814.

[14] Jason E. Chung et al. “A Fully Automated Approach to Spike Sorting.” In: Neuron 95.6 (Sept. 2017). PMCID: PMC5743236, 1381–1394.e6. issn: 0896-6273. doi: 10.1016/j.neuron.2017.08.030. url: 10.1016/j.neuron.2017.08.030.

[15] Tianqi Chen and Carlos Guestrin. “Xgboost: A scalable tree boosting system.” In: Proceedings of the 22nd acm sigkdd international conference on knowledge discovery and data mining. 2016, pp. 785–794.

[16] Keith B. Hengen et al. “Firing Rate Homeostasis in Visual Cortex of Freely Behaving Rodents.” In: Neuron 80.2 (Oct. 2013). PMCID: PMC3816084, pp. 335–342. issn: 0896-6273. doi: 10.1016/j.neuron.2013.08.038. url: 10.1016/j.neuron.2013.08.038.

[17] Jessica A Cardin, Larry A Palmer, and Diego Contreras. “Stimulus feature selectivity in excitatory and inhibitory neurons in primary visual cortex.” In: Journal of Neuroscience 27.39 (2007), pp. 10333–10344.

[18] Cristopher M Niell and Michael P Stryker. “Highly selective receptive fields in mouse visual cortex.” In: Journal of Neuroscience 28.30 (2008), pp. 7520–7536.

[19] Alexander Mathis et al. “DeepLabCut: markerless pose estimation of user-defined body parts with deep learning.” In: Nature Neuroscience 21.9 (Aug. 2018), pp. 1281–1289. issn: 1546-1726. doi: 10.1038/s41593-018-0209-y. url: 10.1038/s41593-018-0209-y.

[20] Keith B. Hengen et al. “Neuronal Firing Rate Homeostasis Is Inhibited by Sleep and Promoted by Wake.” In: Cell 165.1 (Mar. 2016). PMCID: PMC4809041, pp. 180–191. issn: 0092-8674. doi: 10.1016/j.cell.2016.01.046. url: 10.1016/j.cell.2016.01.046.

[21] R Lalonde, HD Kim, and K Fukuchi. “Exploratory activity, anxiety, and motor coordination in bigenic APPswe+ PS1/ΔE9 mice.” In: Neuroscience letters 369.2 (2004), pp. 156–161.

[22] Scott J Webster, Adam D Bachstetter, and Linda J Van Eldik. “Comprehensive behavioral characterization of an APP/PS-1 double knock-in mouse model of Alzheimer’s disease.” In: Alzheimer’s research & therapy 5 (2013), pp. 1–15.

[23] Marc Aurel Busche, et al. “Clusters of hyperactive neurons near amyloid plaques in a mouse model of Alzheimer’s disease.” In: Science 321.5896 (2008), pp. 1686–1689.

[24] Marc Aurel Busche, et al. “Critical role of soluble amyloid-*β* for early hippocampal hyperactivity in a mouse model of Alzheimer’s disease.” In: Proceedings of the National Academy of Sciences 109.22 (2012), pp. 8740–8745.

[25] Jan L. Klee et al. “Reduced firing rates of pyramidal cells in the frontal cortex of APP/PS1 can be restored by acute treatment with levetiracetam.” In: Neurobiology of Aging 96 (Dec. 2020), pp. 79–86. issn: 0197-4580. doi: 10.1016/j.neurobiolaging.2020.08.013. url: 10.1016/j.neurobiolaging.2020.08.013.

[26] Douglas Bates et al. “Fitting linear mixed-effects models using lme4.” In: Journal of statistical software 67 (2015), pp. 1–48.

[27] Anish Mitra, et al. “Human cortical–hippocampal dialogue in wake and slow-wave sleep.” In: Proceedings of the National Academy of Sciences 113.44 (2016), E6868–E6876.

[28] Anish Mitra, et al. “Spontaneous infra-slow brain activity has unique spatiotemporal dynamics and laminar structure.” In: Neuron 98.2 (2018), pp. 297–305.

[29] Disha Shah, et al. “Spatial reversal learning defect coincides with hypersynchronous telencephalic BOLD functional connectivity in APPNL-F/NL-F knock-in mice.” In: Scientific reports 8.1 (2018), p. 6264.

[30] Yasuo Kawaguchi. “Selective cholinergic modulation of cortical GABAergic cell subtypes.” In: Journal of neurophysiology 78.3 (1997), pp. 1743–1747.

[31] Sergio Arroyo et al. “Prolonged disynaptic inhibition in the cortex mediated by slow, non-*α*7 nicotinic excitation of a specific subset of cortical interneurons.” In: Journal of Neuroscience 32.11 (2012), pp. 3859–3864.

[32] Seung-Hee Lee and Yang Dan. “Neuromodulation of brain states.” In: neuron 76.1 (2012), pp. 209–222.

[33] Gabrielle Girardeau, et al. “Selective suppression of hippocampal ripples impairs spatial memory.” In: Nature neuroscience 12.10 (2009), pp. 1222–1223.

[34] Antonio Fernández-Ruiz, et al. “Long-duration hippocampal sharp wave ripples improve memory.” In: Science 364.6445 (2019), pp. 1082–1086.

[35] Raphael Kaplan, et al. “Hippocampal sharp-wave ripples influence selective activation of the default mode network.” In: Current Biology 26.5 (2016), pp. 686–691.

[36] Mattias P Karlsson and Loren M Frank. “Awake replay of remote experiences in the hippocampus.” In: Nature neuroscience 12.7 (2009), pp. 913–918.

[37] E Zhurakovskaya, et al. “Impaired hippocampal-cortical coupling but preserved local synchrony during sleep in APP/PS1 mice modeling Alzheimer’s disease.” In: Scientific reports 9.1 (2019), p. 5380.

[38] Sarah D Cushing, et al. “Impaired hippocampal-cortical interactions during sleep in a mouse model of Alzheimer’s disease.” In: Current Biology 30.13 (2020), pp. 2588–2601.

[39] Naoki Yamawaki, et al. “Long-range inhibitory intersection of a retrosplenial thalamocortical circuit by apical tuft-targeting CA1 neurons.” In: Nature neuroscience 22.4 (2019), pp. 618–626.

[40] Ashley N Opalka, et al. “Hippocampal ripple coordinates retrosplenial inhibitory neurons during slow-wave sleep.” In: Cell Reports 30.2 (2020), pp. 432–441.

[41] Pedro Nascimento Alves, et al. “An improved neuroanatomical model of the default-mode network reconciles previous neuroimaging and neuropathological findings.” In: Communications biology 2.1 (2019), p. 370.

[42] João D Semedo, et al. “Cortical areas interact through a communication subspace.” In: Neuron 102.1 (2019), pp. 249–259.

[43] Srinivas Gorur-Shandilya, et al. “Mapping circuit dynamics during function and dysfunction.” In: Elife 11 (2022), e76579.

[44] Christoph Kapfer, et al. “Supralinear increase of recurrent inhibition during sparse activity in the somatosensory cortex.” In: Nature neuroscience 10.6 (2007), pp. 743–753.

[45] Masuo Ohno, et al. “BACE1 gene deletion prevents neuron loss and memory deficits in 5XFAD APP/PS1 transgenic mice.” In: Neurobiology of disease 26.1 (2007), pp. 134–145.

[46] Mei-Hong Lu, et al. “Inhibition of sphingomyelin synthase 1 ameliorates alzheimer-like pathology in APP/PS1 transgenic mice through promoting lysosomal degradation of BACE1.” In: Experimental neurology 311 (2019), pp. 67–79.

[47] Dan Xia et al. “Novel App knock-in mouse model shows key features of amyloid pathology and reveals profound metabolic dysregulation of microglia.” In: Molecular Neurodegeneration 17.1 (June 2022). PMCID: PMC9188195. issn: 1750-1326. doi: 10.1186/s13024-022-00547-7. url: 10.1186/s13024-022-00547-7.

[48] Rimante Minkeviciene et al. “Amyloid *β*-induced neuronal hyperexcitability triggers progressive epilepsy.” In: Journal of Neuroscience 29.11 (2009), pp. 3453–3462.

[49] Daniel Zarhin, et al. “Disrupted neural correlates of anesthesia and sleep reveal early circuit dysfunctions in Alzheimer models.” In: Cell reports 38.3 (2022).

[50] Heng Zhou, et al. “Disruption of hippocampal neuronal circuit function depends upon behavioral state in the APP/PS1 mouse model of Alzheimer’s disease.” In: Scientific Reports 12.1 (2022), p. 21022.

[51] Jorge J. Palop and Lennart Mucke. “Network abnormalities and interneuron dysfunction in Alzheimer disease.” In: Nature Reviews Neuroscience 17.12 (Nov. 2016). PMCID: PMC8162106, pp. 777–792. issn: 1471-0048. doi: 10.1038/nrn.2016.141. url: 10.1038/nrn.2016.141.

[52] Björn Rasch and Jan Born. “About sleep’s role in memory.” In: Physiological reviews (2013).

[53] James L McClelland, Bruce L McNaughton, and Randall C O’Reilly. “Why there are complementary learning systems in the hippocampus and neocortex: insights from the successes and failures of connectionist models of learning and memory.” In: Psychological review 102.3 (1995), p. 419.

[54] Yo-El S. Ju, Brendan P. Lucey, and David M. Holtzman. “Sleep and Alzheimer disease pathology—a bidirectional relationship.” In: Nature Reviews Neurology 10.2 (Dec. 2013). PMCID: PMC3979317, pp. 115–119. issn: 1759-4766. doi: 10.1038/nrneurol.2013.269. url: 10.1038/nrneurol.2013.269.

[55] Jason D Warren, et al. “Molecular nexopathies: a new paradigm of neurodegenerative disease.” In: Trends in neurosciences 36.10 (2013), pp. 561–569.

[56] Miranda M Lim, Jason R Gerstner, and David M Holtzman. “The sleep–wake cycle and Alzheimer’s disease: what do we know?” In: Neurodegenerative disease management 4.5 (2014), pp. 351–362.

[57] Kathleen Bokenberger et al. “Shift work and risk of incident dementia: a study of two population-based cohorts.” In: European journal of epidemiology 33 (2018), pp. 977–987.

[58] Jee Hoon Roh, et al. “Disruption of the sleep-wake cycle and diurnal fluctuation of *β*-amyloid in mice with Alzheimer’s disease pathology.” In: Science translational medicine 4.150 (2012), 150ra122–150ra122.

[59] David F. Parks et al. “A nonoscillatory, millisecond-scale embedding of brain state provides insight into behavior.” In: Nature Neuroscience (July 2024). issn: 1546-1726. doi: 10.1038/s41593-024-01715-2. url: 10.1038/s41593-024-01715-2.

[60] Zhengyu Ma et al. “Cortical Circuit Dynamics Are Homeostatically Tuned to Criticality In Vivo.” In: Neuron 104.4 (Nov. 2019). PMCID: PMC6934140, 655–664.e4. issn: 0896-6273. doi: 10.1016/j.neuron.2019.08.031. url: 10.1016/j.neuron.2019.08.031.

[61] Eric H Chang, et al. “AMPA receptor downscaling at the onset of Alzheimer’s disease pathology in double knockin mice.” In: Proceedings of the National Academy of Sciences 103.9 (2006), pp. 3410–3415.

[62] Shreaya Chakroborty et al. “Nitric oxide signaling is recruited as a compensatory mechanism for sustaining synaptic plasticity in Alzheimer’s disease mice.” In: Journal of Neuroscience 35.17 (2015), pp. 6893–6902.

[63] Demetris K Roumis and Loren M Frank. “Hippocampal sharp-wave ripples in waking and sleeping states.” In: Current opinion in neurobiology 35 (2015), pp. 6–12.

[64] Jonathan Witton et al. “Disrupted hippocampal sharp-wave ripple-associated spike dynamics in a transgenic mouse model of dementia.” In: The Journal of physiology 594.16 (2016), pp. 4615–4630.

[65] Bartosz Jura, et al. “Deficit in hippocampal ripples does not preclude spatial memory formation in APP/PS1 mice.” In: Scientific reports 9.1 (2019), p. 20129.

[66] Karola Kaefer, et al. “Replay, the default mode network and the cascaded memory systems model.” In: Nature Reviews Neuroscience 23.10 (2022), pp. 628–640.

[67] David T Jones, et al. “Cascading network failure across the Alzheimer’s disease spectrum.” In: Brain 139.2 (2016), pp. 547–562.

[68] H. Braak and E. Braak. “Neuropathological stageing of Alzheimer-related changes.” In: Acta Neuropathologica 82.4 (Sept. 1991), pp. 239–259. issn: 1432-0533. doi: 10.1007/bf00308809. url: 10.1007/BF00308809.

[69] Xiaoyin Chen, et al. “Whole-cortex in situ sequencing reveals peripheral input-dependent cell type-defined area identity.” In: bioRxiv (2022), pp. 2022–11.

[70] Kaoru Yamamoto et al. “Chronic Optogenetic Activation Augments A*β* Pathology in a Mouse Model of Alzheimer Disease.” In: Cell Reports 11.6 (May 2015), pp. 859–865. issn: 2211-1247. doi: 10.1016/j.celrep.2015.04.017. url: 10.1016/j.celrep. 2015.04.017.

[71] Joanna L Jankowsky, et al. “Mutant presenilins specifically elevate the levels of the 42 residue *β*-amyloid peptide in vivo: evidence for augmentation of a 42-specific *γ* secretase.” In: Human molecular genetics 13.2 (2004), pp. 159–170.

[72] Joshua H Siegle et al. “Open Ephys: an open-source, plugin-based platform for multichannel electrophysiology.” In: Journal of neural engineering 14.4 (2017), p. 045003.

[73] Alessio P Buccino, et al. “SpikeInterface, a unified framework for spike sorting.” In: Elife 9 (2020), e61834.

[74] Brendon O Watson, et al. “Network homeostasis and state dynamics of neocortical sleep.” In: Neuron 90.4 (2016), pp. 839–852.

[75] Anli A Liu, et al. “A consensus statement on detection of hippocampal sharp wave ripples and differentiation from other fast oscillations.” In: Nature communications 13.1 (2022), p. 6000.

